# Fic-mediated AMPylation tempers the Unfolded Protein Response during physiological stress

**DOI:** 10.1101/2022.04.27.489443

**Authors:** Amanda K. Casey, Hillery F. Gray, Suneeta Chimalapati, Genaro Hernandez, Andrew Moehlman, Nathan Stewart, Hazel A. Fields, Burak Gulen, Kelly A. Servage, Karoliina Stefanius, Aubrie Blevins, Elena Daoud, Bret Evers, Helmut Krämer, Kim Orth

## Abstract

The proper balance of synthesis, folding, modification and degradation of proteins, also known as protein homeostasis, is vital to cellular health and function. The unfolded protein response (UPR) is activated when the mechanisms maintaining protein homeostasis in the endoplasmic reticulum (ER) become overwhelmed. However, prolonged or strong UPR responses can result in elevated inflammation and cellular damage. Previously, we discovered that the bifunctional enzyme Fic can modulate the UPR response via post-translational modification of BiP by AMPylation and deAMPylation. Loss of *fic* in *Drosophila* leads to vision defects and altered UPR activation in the fly eye. To investigate the importance of Fic-mediated AMPylation in a mammalian system, we generated a conditional null allele of *Fic* in mice and characterized the effect of *Fic* loss on the exocrine pancreas. Compared to controls, *Fic^-/-^* mice exhibit elevated serum markers for pancreatic dysfunction and display enhanced UPR signaling in the exocrine pancreas in response to physiologic and pharmacological stress. In addition, both *fic^-/-^* flies and *Fic^-/-^* mice show reduced capacity to recover from damage by stress that triggers the UPR. These findings show that Fic- mediated AMPylation acts as a molecular rheostat that is required to temper the UPR response in the mammalian pancreas during physiological stress.

## Introduction

Protein homeostasis is regulated by proper synthesis, folding, modification and degradation of proteins and is vital to cellular health. In the endoplasmic reticulum (ER), when the load of unfolded proteins is excessive, the unfolded protein response (UPR) is activated, triggering signaling pathways that result in changes to protein synthesis, modification, and degradation, until the load of unfolded proteins is resolved. If the burden of unfolded proteins is prolonged and/or remains high, pro-apoptotic pathways can be activated. The activation of the UPR is, in part, regulated by the Hsp70 protein chaperone BiP, a protein that binds and helps fold proteins as they pass through the ER checkpoint and into the secretory pathway. Depending on the level of unfolded protein, complex signaling networks are activated and respond in accordance to the severity. Mild to moderate levels of UPR signaling are promote with cell recovery and cell survival whereas strong and prolonged UPR signaling lead to apoptosis (1, 2). These responses are mediated by three distinct ER signaling branches: Inositol-requiring enzyme-1α (IRE1α); protein kinase R-like ER kinase (PERK); and activating transcription factor 6α (ATF6α) (3).

In addition, the UPR stress can be divided into two phases: the *adaptive phase* and the *maladaptive phase* (4, 5). For the adaptive phase, the UPR induction responds to mild to moderate stress and promotes pro-survival and restorative mechanisms to promote ER homeostasis (4). By contrast, the maladaptive phase is induced by chronic and severe ER stress resulting in the activation of pro-inflammatory responses and apoptosis (4, 6). Disruption of ER homeostasis is predicted to play a key role in the integrated stress response (ISR) and the progression of many neurodegenerative, inflammatory, and metabolic disorders (7–9). Elucidating the roles that the UPR plays in modulating ER stress provides potential therapeutic targets to treat or prevent the death of the cells subjected to prolonged ER stress and ameliorate UPR-related degenerative diseases.

Previously, the activity of BiP was shown to be regulated by a post-translational modification (PTM) called AMPylation (10). AMPylation is a reversible PTM best described as the covalent linkage of adenosine monophosphate (AMP) to the hydroxyl group of a serine, threonine, or tyrosine residue (11). Initially discovered in the 1960’s with a nucleotidyl transferase domain (12), protein AMPylation was rediscovered in 2009 with a Fic domain from a bacterial pathogen that is also conserved in eukaryotic organisms (13, 14). To date, only two AMPylating enzymes have been identified in metazoans: Fic (also known as FicD and HYPE) localizes in the ER, and SelO localizes in the mitochondria (15, 16).

Using *Drosophila* as an animal model, we found that Fic is responsible for reversible AMPylation of BiP (17). During low ER stress or resting cells, Fic AMPylates BiP, thereby creating an inactive pool of BiP in the ER lumen (17, 18). When ER stress rises, BiP is deAMPylated and returned to an active state (10). Since this discovery, other labs have confirmed that this function is conserved in other metazoans, including *C. elegans*, rodents, and humans (16, 19, 20). We and others then demonstrated that Fic has dual catalytic activity for both the AMPylation and deAMPylation of metazoan BiP and this activity changes depending on levels of ER stress (21, 22)

Further studies on the Drosophila model revealed that Fic plays a crucial role in protein homeostasis for metazoans. For *Fic* null flies (*fic^-/-^*), the gross morphology of the fly eye appeared normal, albeit they exhibited mild vision defects. When acute physiological ER stress was induced in fly eyes by exposure to continuous light, photoreceptors in wild-type flies, but not in *fic^-/-^* mutants, could adapt (23). The damaged *fic^-/-^* eyes exhibited severe structural defects in rhabdomeres (rhodopsin containing membranes), elevated IRE1 activity, and reduced neurotransmission (23). Flies expressing *BiP^S366A^* that are unable to be AMPylated at Ser366 phenocopied the *fic^-/-^* flies, with damaged rhabdomeres and loss of postsynaptic responses for photoreceptors stressed with continuous light. Taken together, these studies support the proposal that having a reserve of inactive AMPylated BiP that can be immediately accessed by deAMPylation, allows cells to more efficiently deal with physiological stress.

Overall, we propose Fic acts as a rheostat that tempers the cellular response to stress and maintains homeostasis by deAMPylating a reserve pool of modified BiP, thereby increasing levels of active BiP to alleviate mild ER stress. When the rheostat is disrupted, either by the absence of Fic or by a mutation in BiP that hampers its AMPylation, recovery from physiologically stressed cells is hindered as there is no resource for immediate access to additional BiP pools. In the absence of this pool, more BiP can only be provided by the time consuming UPR-induced transcription and translation of de novo BiP, coincidently with the triggering of ISR.

Based on these findings, we predicted that Fic is also required for the proper regulation of physiological stress in mammals. To address this hypothesis, we generated a conditional knockout line of Fic in the mouse. As with flies, the *Fic^-/-^* animals are viable, fertile, and appear healthy upon initial inspection. However, closer characterization of *Fic^-/-^* pancreata revealed altered responses to physiological and pathological stresses, with significant changes in UPR-induced signaling. We hypothesize that without Fic, the balance and threshold between the *adaptive phase* and the *maladaptive phase* of the UPR is shifted in tissues that rely heavily on ER secretory pathway to maintain protein homeostasis. Interestingly, we observe marked resilience in wild-type flies and mice when dealing with repeated stress. By contrast, both *fic^-/-^* flies and *Fic^-/-^* mice lack the ability to efficiently recover from these stresses resulting in damaged eyes and scarred pancreas, respectively. Taken together, our findings support the hypothesis that metazoan Fic plays a critical role by acting as a rheostat for the regulation of the UPR and protein homeostasis, likely to be important for the resilience of terminally differentiated, professional secretory cells that must respond to fluctuating needs of an organ.

## Results

### Conditional Deletion of FicD

To determine the role of Fic-mediated AMPylation in the mammalian system, we chose to generated a conditional knockout line of *Fic* in the mouse. Using CRISPR-Cas9 technology, we generated a floxed allele of *Fic* (*Fic^fl^*) in which 2 LoxP sites were integrated upstream of and within exon 3 (**Figure 1A**). In addition, a single FLAG epitope was inserted into the C-terminal sequence of Fic’s coding sequence (**Figure 1A**, **Figure 1 – figure supplement 1).** Expression of Cre leads to Cre-Lox recombination that results in a non-functional Fic gene (Fic^-^) that is deleted for both the TPR and Fic domains that are required for the targeting and AMPylation of BiP, respectively. The remaining truncated gene encodes only a small N-terminal peptide with the transmembrane sequence (**Figure 1A**). *Fic^fl^* mice were bred to CAG-Cre transgenic mice (24) to generate *Fic^+/-^* mice which were then backcrossed with *Fic^fl^* mice to generate *Fic^fl/-^* mice. Germline transmission of this allele was confirmed through PCR and sequencing (**Figure 1 – figure supplement 2**). *Fic^fl/-^* mice were intercrossed to obtain sibling cohorts of *Fic^fl/fl^* and *Fic^-/-^* mice that were subsequently used in this study. *Fic^-/-^* mice were indistinguishable compared to *Fic^fl/fl^* and *Fic^fl/-^* littermates in viability, appearance, and weight (**Figure 1 – figure supplement 3**).

**Figure 1.**
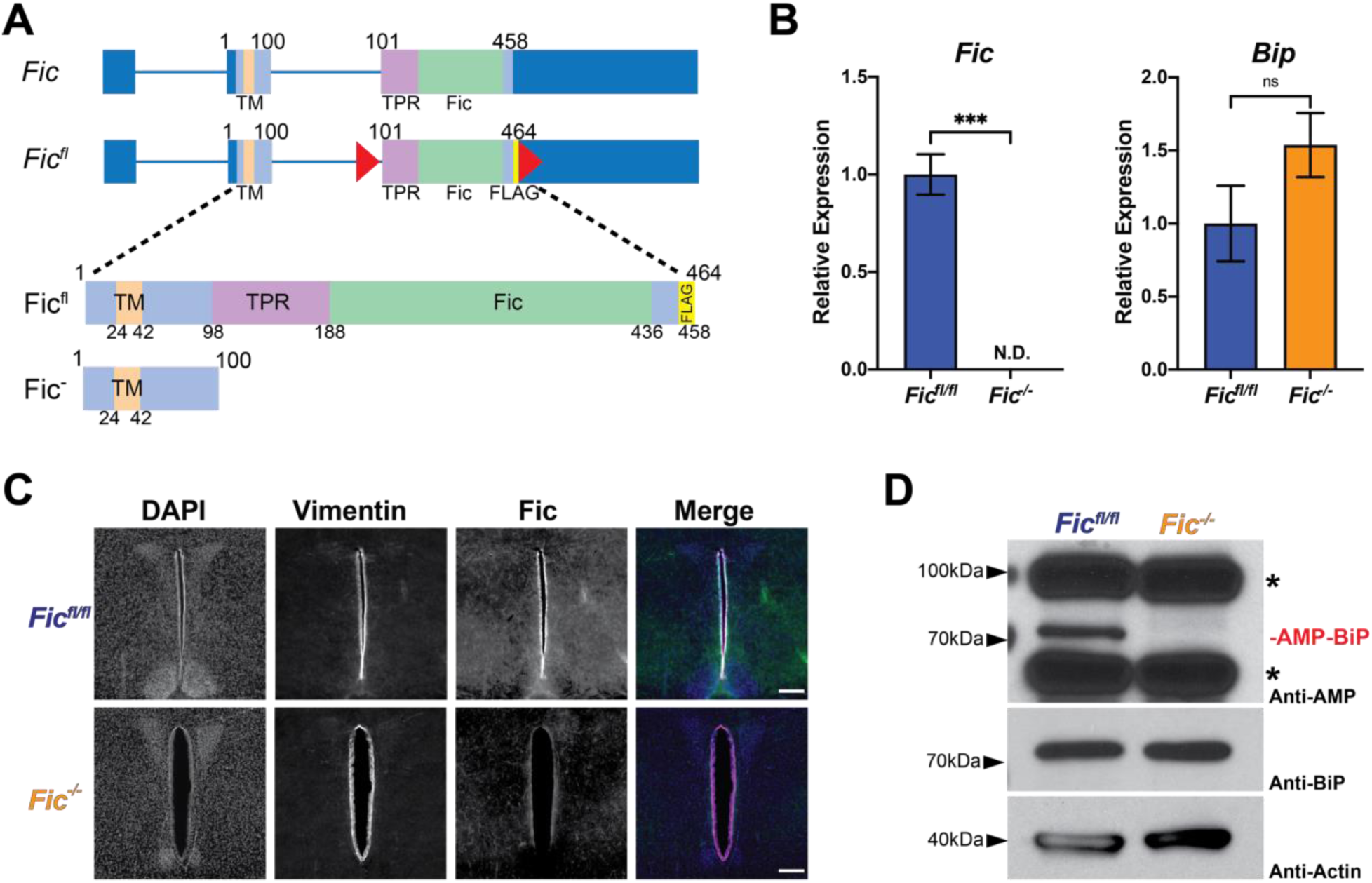
Conditional Deletion of FicD. A) A schematic representation of wildtype and floxed allele of Fic (*Fic^fl^*) in which LoxP sites were inserted into intron 2 and Exon 3 of the Fic gene. A 6 amino acid FLAG sequence was added to the C-terminus of the Fic ORF. B) Quantification of *Fic* and *BiP* mRNA analyzed by qPCR from *Fic^fl/fl^* (blue bar) and *Fic^-/-^* (orange bar) mouse liver after 12 hours fasting. N=3. Bars indicate mean relative expression and error bars represent standard error. Fic mRNA was below detection cutoff in *Fic^-/-^* samples. Statistics were performed using GraphPad Prism 9 using an unpaired t-test. N.D., not detected; ***, p < 0.001; ns, not significant. C) Representative image of Fic and Vimentin immunohistochemistry in coronal section of murine third ventricle. Scale bar, 200μM D) Representative western blot of liver lysates isolated from *Fic^fl/fl^* and *Fic^-/-^* mice. Blots were probed with anti-AMP, anti-BiP, and anti-Actin antibodies. Asterisks indicate reactivity to AMP antibody unrelated to Fic expression.

### Fic^-/-^ mice lack Fic protein and RNA transcript and BiP AMPylation

Using qPCR analysis of liver cDNA we confirmed loss of *Fic* transcript corresponding to exon 3 of Fic, validating the Fic knockout (**Figure 1B**). Levels of BiP transcript were not significantly altered in *Fic^fl/fl^* and *Fic^-/-^*liver. We then attempted to validate deletion of *Fic* in mice using Western blot analysis of lysates from various tissues; however, due to the low expression level of endogenous protein, we were unable to detect Fic protein in wild-type or *Fic^fl/fl^* controls (data not shown). As an alternative, we used immunofluorescence in various tissues known to express Fic and were able to detect a signal for Fic in coronal brain sections, strikingly in tanycytes in the third ventricle of the hypothalamus (**Figure 1C, Figure 1 – figure supplement 4**). In corresponding sections from *Fic^-/-^* mice, we were unable to detect a signal for Fic protein, supporting the genetic and mRNA expression data that Fic is not expressed in *Fic^-/-^* mice.

We next analyzed tissue for the presence or absence of Fic-mediated BiP AMPylation. Whole cell lysates of livers from *Fic^fl/fl^* and *Fic^-/-^* littermates were analyzed for the presence of AMPylated BiP. Both Western-blot analysis (**Figure 1D**) and mass spectrometry analysis (**Figure 1 – figure supplement 5**) indicate that BiP is no longer AMPylated in *Fic^-/-^* mice.

### Fasted Fic^-/-^ mice display elevated serum Amylase levels

We predicted that *Fic^-/-^* mice would have dysfunction in tissues that rely heavily on UPR to maintain proteostasis. Although many tissues are known to require this regulation, we decided to focus our initial study on Fic in the pancreas, a tissue that is well documented to rely on the UPR for proper exocrine and endocrine function (25, 26). Using the UPR fasting model, we screened *Fic^fl/fl^* and *Fic^-/-^ mice* for serum markers that might indicate abnormalities in pancreatic function. A cohort of 10-11 week old male mice were fasted overnight (∼14 hours) with unrestricted access to water before sacrifice. Compared to *Fic^fl/fl^* and *Fic^fl/-^* controls, fic null mice have normal weights (**Figure 1 – figure supplement 3**) and appear generally healthy. However, fasted serum levels of Amylase were found to be significantly elevated in our *Fic^-/-^* cohorts (**Figure 2 – figure supplement 1**). Of note, serum lipase levels and fasting glucose levels were not affected in our *Fic^fl/fl^* and *Fic^-/-^* mice. Since high Amylase has been linked to hepatic disfunction(27), we looked at another serum marker for hepatic disfunction, aspartate aminotransferase (AST), to see if levels had changed, but levels of AST appeared normal in both *Fic^fl/fl^* and *Fic^-/-^* mice (**Figure 2 – figure supplement 1**). Histopathological analysis of both pancreas and liver revealed no detectable defect in either tissue (**Figure 2 – figure supplement 2**).

### Physiological stressed fasted-fed Fic^-/-^ mice display altered UPR signaling in the pancreas

Based on the observed changes in serum amylase in our *Fic^-/-^* mice, we predicted that *Fic^-/-^* mice would show changes in UPR activation in the exocrine pancreatic tissue. To determine if this was the case, we utilized a mild physiological stress of fasting-feeding to activate the UPR in the exocrine pancreas. Fasting-feeding is a well-established method used to activate mild UPR in pancreas through the stimulation of pancreatic digestive enzyme synthesis (28, 29). In this study, sibling cohorts of 6-8 week old male mice were split into three groups: fasted (for 16 hours), fasted-fed (fasted 14-hours, fed 2 hours), and fasted-fed-recovery (fasted 12 hours, fed 2 hours, fasted 2 hours) (**Figure 2A**).

**Figure 2.**
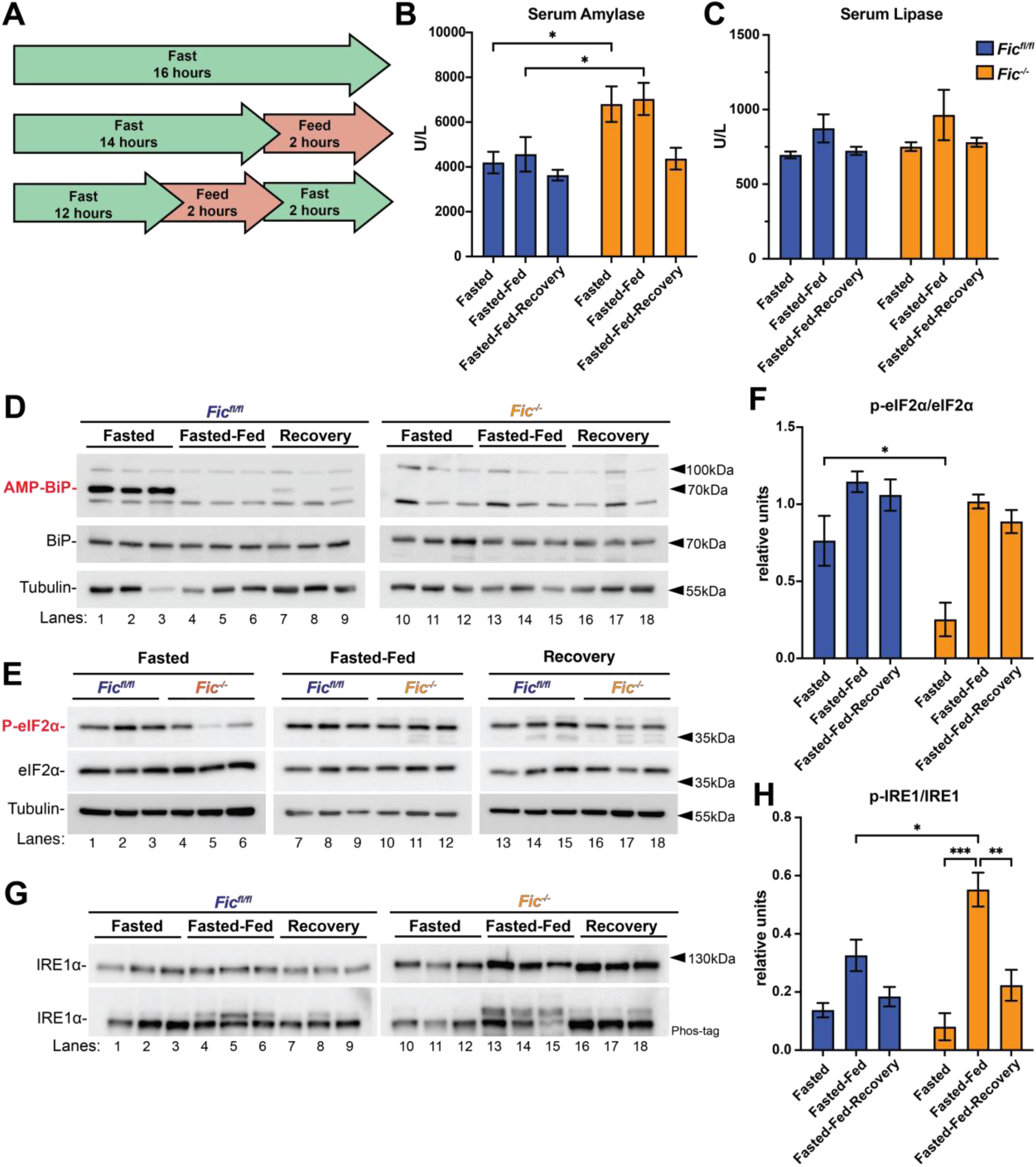
Response of fasted-fed Fic^fl/fl^ and Fic^-/-^ pancreata. A) A schematic representation of fasted, fasted-fed, and fasted-fed-recovery experimental conditions. B) Quantification of serum Amylase in *Fic^fl/fl^ and Fic^-/-^* mice under fasted, fasted-fed, and fasted-fed-recovery condions. Statistics were performed using GraphPad Prism 9 using an 2-way ANOVA. N=8. *, p < 0.05. Bars indicate mean and error bars represent standard error. C) Quantification of serum Lipase in *Fic^fl/fl^ and Fic^-/-^* mice under fasted, fasted-fed, and fasted-fed-recovery condions. N=8. Bars indicate mean and error bars represent standard error. D) Representative western blot of P2 lysate fractions from isolated from *Fic^fl/fl^* and *Fic^-/-^* pancreas. Blots were probed with anti-AMP, anti-BiP, and anti-Tubulin antibodies. E) Representative western blot of S2 lysate fractions from isolated from *Fic^fl/fl^* and *Fic^-/-^* pancreas. Blots were probed with anti-phospho-eiF2α, anti- eiF2α, and anti-Tubulin antibodies. F) Quantification of relative anti-phospho-eiF2α in E. Statistics were performed using GraphPad Prism 9 using 2-way ANOVA. N=3. *, p < 0.05 G) Representative western blot of P2 lysate fractions from isolated from *Fic^fl/fl^* and *Fic^-/-^* pancreas. Blots were probed with anti-Ire1 antibodies. H) Quantification of relative phospho Ire1 in (G) Statistics were performed using GraphPad Prism 9 using an 2-way ANOVA. N=3. *, p < 0.05; **, p < 0.01; ***, p < 0.001.

Compared to *Fic^fl/fl^* controls, elevated serum amylase levels, but not lipase, were detected in fasted and fasted-fed groups of *Fic^-/-^* mice (**Figure 2B-C**). Notably, serum amylase levels were not increased in *Fic^-/-^* mice in the fasted-fed-recovery group. Western analysis of pancreatic lysates in *Fic^fl/fl^* samples from a separate cohort (6-8 week old female mice) confirmed that BiP AMPylation is present during fasted conditions and lost during fasted-fed conditions. Of note, AMPylation of BiP returns in the fasted-fed-recovery group, albeit reduced (**Figure 2D**). Western analysis of phospo-eIF2α and eIF2α showed reduced eIF2α phosphorylation in fasted *Fic^-/-^* mice whereas phospho-eIF2α was not found to differ during fasted-fed and fasted-fed-recovery conditions (**Figure 2E-F**). Western analysis performed on Phos-tag gels indicated that activation of IRE1 was observed to be stronger in fasted-fed *Fic^-/-^* mice compared to *Fic^fl/fl^* controls (**Figure 2G-H**).

To further assess the activation of the UPR in the pancreas, levels of UPR induced transcripts were analyzed using qualitative PCR (qPCR). As previously reported (29), feeding after fasting resulted in a significant increase in UPR-regulated transcripts in mice (**Figure 3, Figure 3 – figure supplement 1**). In *Fic^fl/fl^* mice, levels of *Fic* were found to increase during fasted-fed conditions and then diminish during fasted-fed-recovery (**Figure 3 – figure supplement 1**). Notably, in *Fic^-/-^* mice the many UPR-induced transcripts were elevated compared to *Fic^fl/fl^* controls. Levels of *BiP, Atf4, Xbp1s,* and *Chop* were all found to be significantly increased in *Fic^-/-^* mice during fasted-fed conditions (**Figure 3A-3D**).

**Figure 3.**
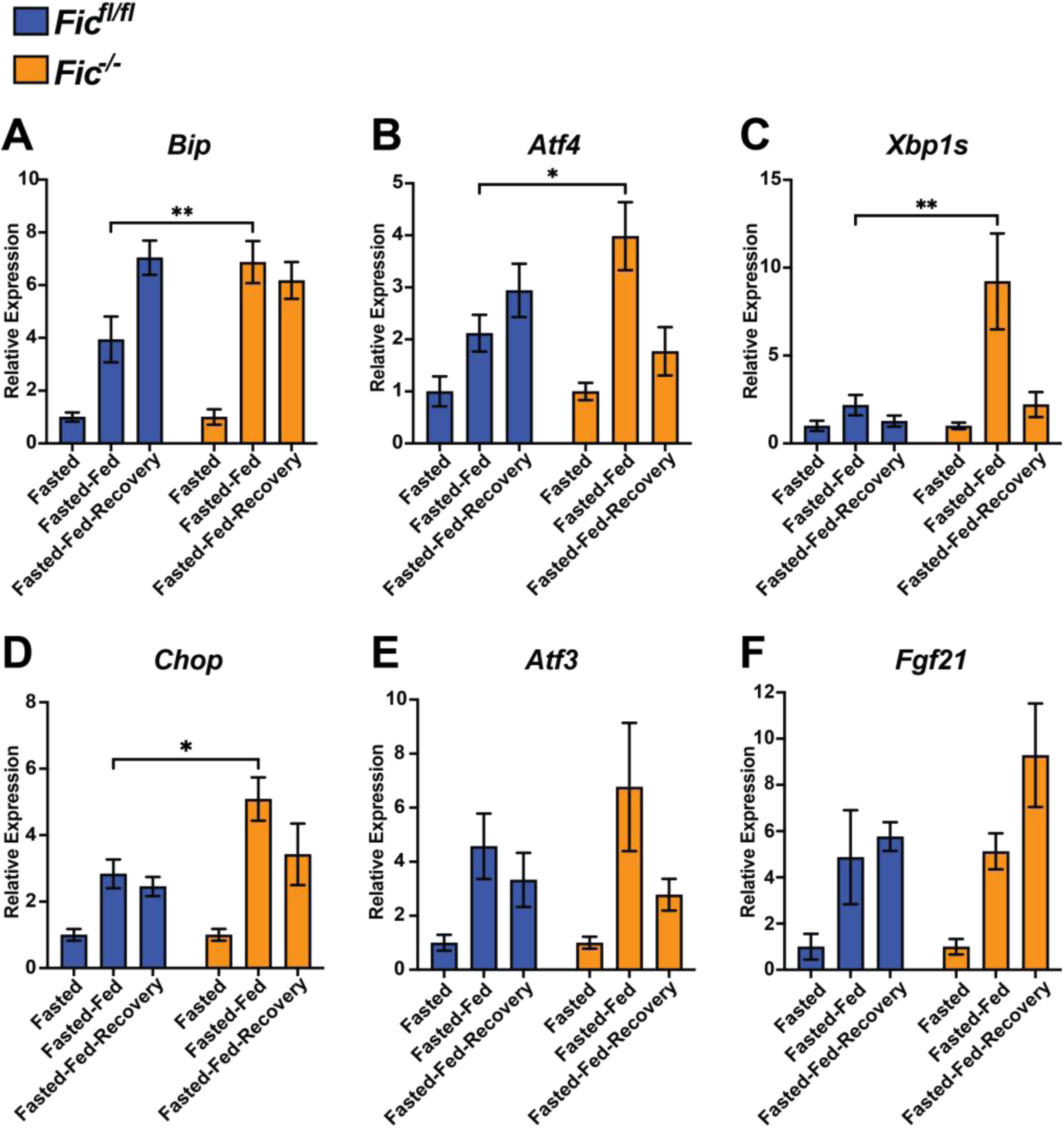
UPR signaling in fasted-fed Fic^fl/fl^ and Fic^-/-^ pancreata. A-F) Quantification of *BiP, ATF4, Xbp1s, Chop, ATF3,* and *Fgf21* mRNA analyzed by qPCR from *Fic^fl/fl^* (blue bar) and *Fic^-/-^* (orange bar) mouse pancreas after fasting, fast-feeding, and fast-feed-recovery. Expression values were normalized to that of the housekeeping gene *U36B4.* Bars indicate mean relative expression compared to fasted controls, and error bars represent standard error. Statistics were performed using GraphPad Prism 9 using an 2-way ANOVA. N=8. *, p < 0.05; **, p < 0.01.

However, not all UPR responsive genes followed this pattern. Changes in levels of *Atf3* and *Fgf21* were not found to differ significantly between *Fic^fl/fl^* and *Fic^-/-^* mice upon fast-feeding (**Figure 3E-F**). Of note, no significant differences of these transcript levels were observed in the fasted-fed-recovery group, signifying this difference in UPR induction is short lived.

### Stressing Fic^-/-^ mice with pancreatic Caerulein treatment reveals changes in UPR response

Next, we wanted to assess the fitness of *Fic^-/-^* mice in response to a maladaptive, pathological ER stress and injury to the pancreas. For this we used a well-established model for pancreatic injury by injecting mice with caerulein, a cholecystokinin (CCK) analog and secretagogue (30, 31). In this study, pancreatitis was induced in 8-10 week old female cohorts with 7 hourly IP injections of saline or caerulein. Animals were sacrificed at 1hr, 4hrs, 8hrs, 24hrs, and 72hrs after the first injection (**Figure 4A**). Immediately upon sacrifice, serum was collected from each mouse and the pancreas was collected for RNA and histopathology.

**Figure 4.**
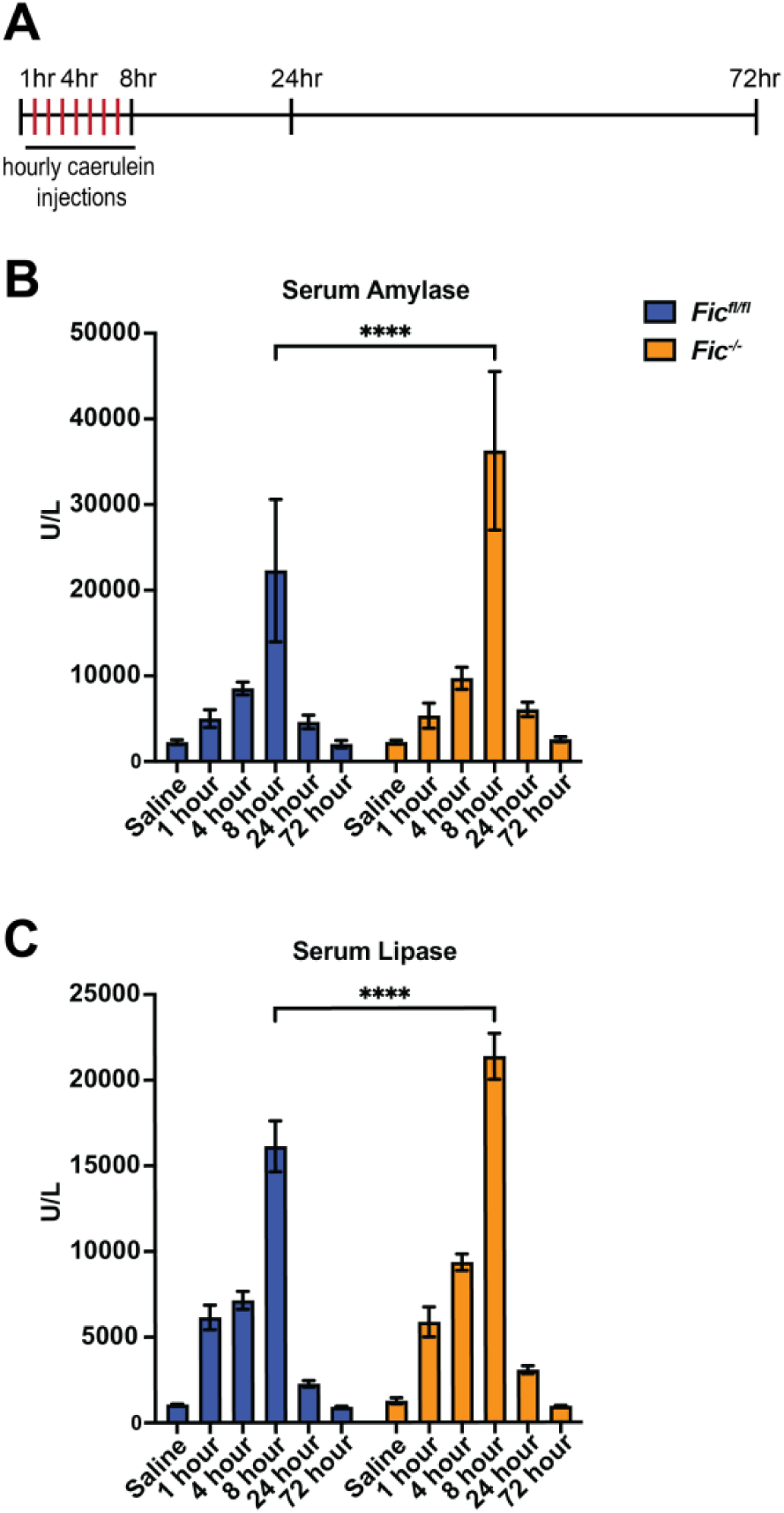
Caerulein-induced pancreatitis in Fic^fl/fl^ and Fic^-/-^ mice. A) A schematic representation of caerulein-induced acute pancreatitis induction. B-C) Quantification of serum Amylase and Lipase in *Fic^fl/fl^ and Fic^-/-^* mice over 72 hours of caerulein-induced acute pancreatitis. Bars indicate mean and error bars represent standard error. Statistics were performed using GraphPad Prism 9 using an 2-way ANOVA. N=5-7. ****, p < 0.0001.

As previously reported (30), treatment with caerulein induces an acute pancreatitis. Analysis of serum revealed elevated levels of Amylase and Lipase in caerulein-treated mice. In addition, levels of serum Amylase and Lipase were significantly increased in *Fic^-/-^* mice compared to *Fic^fl/fl^* at 8 hours after the first injection of caerulein (**Figure 4B-C**).

To assess if caerulein injury differentially alters the UPR response in *Fic^-/-^* mice compared to *Fic^fl/fl^* mice, we assessed the levels of UPR induced transcripts in saline and caerulein-treated mice by qPCR. (**Figure 5**). As previously reported, UPR-induced transcripts are elevated upon caerulein induced injury and resolve over the course of injury recovery (32–35). Whereas both *Fic^fl/fl^* and *Fic^-/-^* mice displayed similar levels of increased UPR transcript induction, the peak of transcript changes appeared earlier in *Fic^-/-^* mice.

**Figure 5.**
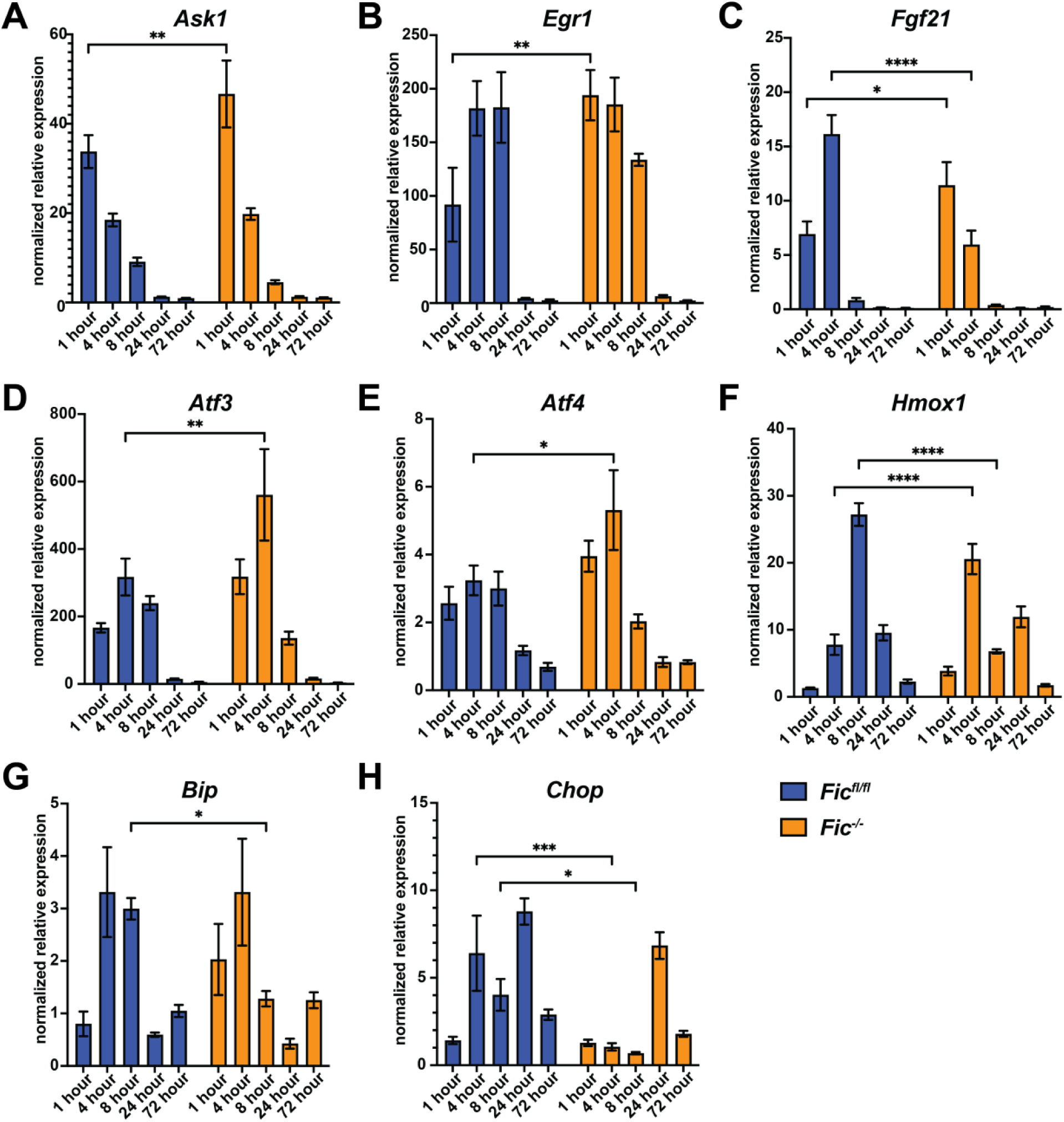
UPR signaling in caerulein-induced acute pancreatitis in fasted-fed Fic^fl/fl^ and Fic^-/-^ mice. A-H) Quantification of *Ask1*, *Egr1*, *Fgf21*, *Atf3*, *Atf4*, *Hmox1*, *BiP*, and *Chop* mRNA analyzed by qPCR from *Fic^fl/fl^* (blue bar) and *Fic^-/-^* (orange bar) mouse pancreas during 72 hours of caerulein-induced acute pancreatitis and recovery. Expression values were normalized to that of the housekeeping gene *U36B4.* Bars indicate mean relative expression compared to Saline-treated controls at each timepoint. Error bars represent standard error. Statistics were performed using GraphPad Prism 9 using an 2-way ANOVA. N=5-7. *, p < 0.05; **, p < 0.01; ***, p < 0.001; ****, p < 0.0001.

One hour after first injection of caerulein, levels of the UPR-induced transcripts *Ask1*, *Egr1*, and *Fgf21* transcript were found to be elevated in *Fic^-/-^* mice compared to wild-type mice (**Figure 5A-C**). However, at 4 hours after the first injection, *Ask1* and *Egr1* levels are comparable in wildtype and *Fic^-/-^* mice and *Fgf21* transcript levels are significantly lower than in wildtype. Similarly, at 4 hours after the first injection, levels of *Atf3*, *Atf4*, and *Hmox1* transcripts are elevated in *Fic^-/-^* mice compared to *Fic^fl/fl^* (**Figure 5D-F**). By 8 hours after first injection, transcript levels of *Atf3* and *Atf4* are comparable between *Fic^fl/fl^* and *Fic^-/-^* mice whereas *Hmox1* transcript levels are significantly lower in *Fic^-/-^* mice. *BiP* transcript levels also drop significantly in *Fic^-/-^* mice at 8 hours **(Figure 5G**). Expression patterns of *Chop* are also significantly altered, with reduced transcript levels for *Fic^-/-^* both at 4 and 8 hours after first caerulein injection (**Figure 5H**). With the exception of *Chop*, UPR transcripts appear to peak and diminish in the pancreas of caerulein treated *Fic^-/-^* mice early compared to *Fic^fl/fl^*.

### Fic is required for recovery of ER stress in both Drosophila and mice

Next, we asked if Fic was required for cellular recovery after prolonged ER stress. In previous studies using *Drosophila* as a model, we found that *fic^-/-^* flies were maladaptive to light stress. In this system, three days of constant light caused damage to ommatidial structures found in the *fic^-/-^* compound eye and enhanced UPR activation (23). In these previous experiments, we found that the defects in rhabdomere integrity were completely reversible in wild-type flies, but, significantly, *fic^-/-^* flies only partially recovered after three days of normal light/dark conditions.

As an extention of these previous experiments, we wanted to ask whether a repetitive version of this stress in flies results in a degenerative effect in affected tissues. To test this, we repeated treatments of constant light (LL) for three days, followed by three days of recovery (LD) on the same cohort of flies and scored for presence or lack of intact deep pseudopupils (DPPs), a visual indicator for disruption of ommatidal structure (**Figure 6A**). As previously observed, for the first round of LL and LD treatment, wild-type flies completely recovered but *fic^-/-^* flies showed only a 91% recovery. We found that with a second round of LL and LD treatment, wild-type flies retained their ability to fully recover but *fic^-/-^* flies had only a 82% recovery. Finally, by the third repeat of this stress, wild-type flies continued to fully recover whereas only 56% of *fic^-/-^* flies retained intact DPPs.

**Figure 6.**
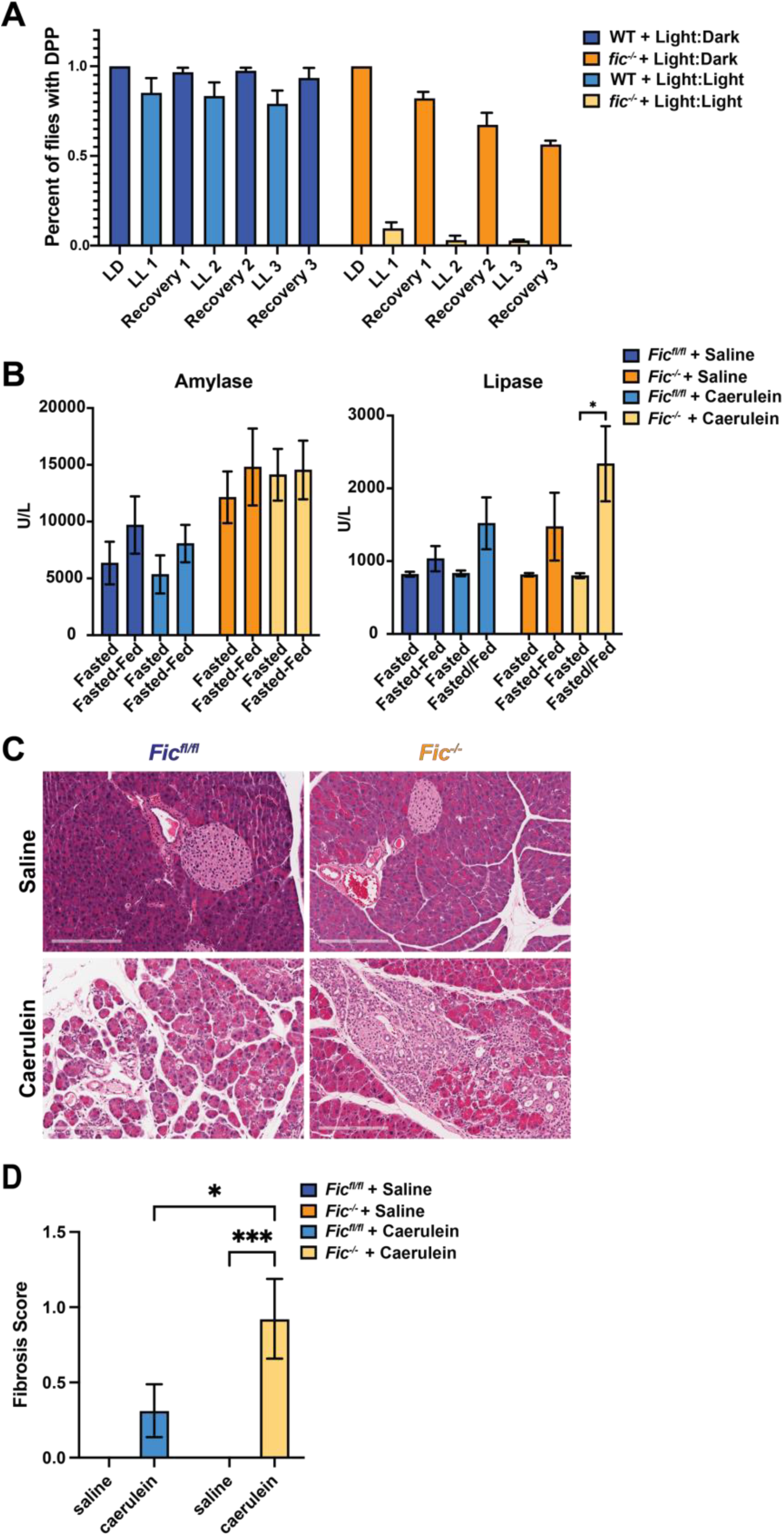
Fic is required for recovery after stress. A) Quantification of flies possessing intact deep pseudopuils (DPPs) at each indicated time. Bars indicate mean, and error bars represent standard error. N=∼150. B) Quantification of serum Amylase and Lipase in *Fic^fl/fl^ and Fic^-/-^* mice 7 days after caerulein-induced acute pancreatitis. Bars indicate mean, and error bars represent standard error. Statistics were performed using GraphPad Prism 9 using an 2-way ANOVA. N=6-8. *, p < 0.05. C) Representative hematoxylin and eosin stained images of pancreas of *Fic^fl/fl^* and *Fic^-/-^* mice 7 days after treatment with saline or caerulein. D) Quantification of pancreas with fibrosis, *Fic^fl/fl^* and *Fic^-/-^* mice 7 days after treatment with saline or caerulein. Bars indicate mean, and error bars represent standard error. Statistics were performed using GraphPad Prism 9 using an 2-way ANOVA. N=13-16 ***, p < 0.001.

Based on the changes in UPR response kinetics and increased serum amylase and lipase levels in caerulein-treated *Fic^-/-^* mice, we hypothesized that *Fic^-/-^* mice might also exhibit more severe and prolonged histopathological changes in the pancreas upon caerulein-induced pancreatitis. The severity of pancreatitis in each saline and caerulein-treated mouse was scored by severity of pancreatic edema, inflammatory infiltrate and necrosis as previously described (36). Based on these criteria, *Fic^fl/fl^* and *Fic^-/-^* mice exhibited comparable gross histological changes to the pancreas with inflammation, edema, and necrosis observed in both cohorts over the course of the 72 hour experiment (**Figure 6 – figure supplement 1-2 and Table 1**), revealing no detectable difference in pancreatitis severity tissue after one round of acute caerluein injury.

**Table 1.** histopathological scoring of caerulein pancreatitis

| GENOTYPE | TREATMENT | Time | Edema | Inflammation | Necrosis | TOTAL |
| --- | --- | --- | --- | --- | --- | --- |
| Fic<sup>fl/fl</sup> | SALINE | 1hr | 0 ± 0 | 0 ± 0 | 0 ± 0 | 0 ± 0 |
| Fic<sup>-/-</sup> | SALINE | 1hr | 0 ± 0 | 0 ± 0 | 0 ± 0 | 0 ± 0 |
| Fic<sup>fl/fl</sup> | CAERULEIN | 1hr | 0 ± 0 | 0 ± 0 | 0 ± 0 | 0 ± 0 |
| Fic<sup>-/-</sup> | CAERULEIN | 1hr | 0 ± 0 | 0 ± 0 | 0 ± 0 | 0 ± 0 |
| Fic<sup>fl/fl</sup> | SALINE | 4hr | 0 ± 0 | 0 ± 0 | 0 ± 0 | 0 ± 0 |
| Fic<sup>-/-</sup> | SALINE | 4hr | 0 ± 0 | 0 ± 0 | 0 ± 0 | 0 ± 0 |
| Fic<sup>fl/fl</sup> | CAERULEIN | 4hr | 1 ± 0 | 0 ± 0 | 0 ± 0 | 0 ± 0 |
| Fic<sup>-/-</sup> | CAERULEIN | 4hr | 1 ± 0 | 0 ± 0 | 0 ± 0 | 0 ± 0 |
| Fic<sup>fl/fl</sup> | SALINE | 8hr | 0 ± 0 | 0 ± 0 | 0 ± 0 | 0 ± 0 |
| Fic<sup>-/-</sup> | SALINE | 8hr | 0 ± 0 | 0 ± 0 | 0 ± 0 | 0 ± 0 |
| Fic<sup>fl/fl</sup> | CAERULEIN | 8hr | 1.83 ± 0.40 | 2.17 ± 0.17 | 2.5 ± 0.22 | 6.5 ± 0.72 |
| Fic<sup>-/-</sup> | CAERULEIN | 8hr | 2.4 ± 0.40 | 2.2 ± 0.37 | 2.6 ± 0.40 | 7.2 ± 1.11 |
| Fic<sup>fl/fl</sup> | SALINE | 24hr | 0 ± 0 | 0 ± 0 | 0 ± 0 | 0 ± 0 |
| Fic<sup>-/-</sup> | SALINE | 24hr | 0 ± 0 | 0 ± 0 | 0 ± 0 | 0 ± 0 |
| Fic<sup>fl/fl</sup> | CAERULEIN | 24hr | 2.25 ± 0.25 | 2.38 ± 0.18 | 2.25 ± 0.16 | 6.88 ± 0.44 |
| Fic<sup>-/-</sup> | CAERULEIN | 24hr | 2.29 ± 0.18 | 2.43 ± 0.20 | 2.14 ± 0.26 | 6.86 ± 0.51 |
| Fic<sup>fl/fl</sup> | SALINE | 72hr | 0 ± 0 | 0 ± 0 | 0 ± 0 | 0 ± 0 |
| Fic<sup>-/-</sup> | SALINE | 72hr | 0.14 ± 0.14 | 0.29 ± 0.29 | 0.29 ± 0.28 | 0.71 ± 0.71 |
| Fic<sup>fl/fl</sup> | CAERULEIN | 72hr | 0.67 ± 0.21 | 2 ± 0.26 | 2 ± 0.26 | 4.67 ± 0.67 |
| Fic<sup>-/-</sup> | CAERULEIN | 72hr | 0.71 ± 0.18 | 1.86 ± 0.34 | 1.71 ± 0.36 | 4.29 ± 0.81 |

average scores+/- S.E.M.

Previous reports indicate wild-type mice typically fully recover from caerulein-induced acute pancreatitis by 7 days post-injury (37). We wanted to assess how *Fic^fl/fl^* and *Fic^-/-^* mice recovered from caerulein-induced pancreatitis. A cohort of 6-8 week old *Fic^fl/fl^* and *Fic^-/-^* male mice was induced for pancreatitis with 7 hourly IP injections of saline or caerulein and left to recover for 7 days. Mice were then split into two groups: a fasted and fasted-fed group as previously described (**Figure 2A**). Immediately upon sacrifice, RNA and tissue for histopathology from the pancreas of each mouse were collected.

As previously described (**Figure 2B**) *Fic^-/-^* mice had elevated serum amylase levels upon fasting and fasting-feeding, but the elevation of serum amylase was no longer affected, indicating that both *Fic^fl/fl^* and *Fic^-/-^* mice had recovered from the initial caerulein-induced injury (**Figure 6B**). Notably, serum Lipase of caerulein-treated *Fic^-/-^* mice after fast-feeding was elevated. Moreover, histopathological scoring of the pancreas showed an increased incidence of fibrosis in caerulein treated *Fic^-/-^* mice than *Fic^fl/fl^* after 7 days of recovery (**Figure 6C-D, Table 2**). This data suggests that *Fic^-/-^* mice have increased scarring and reduced capacity for recovery from tissue damage.

**Table 2.**
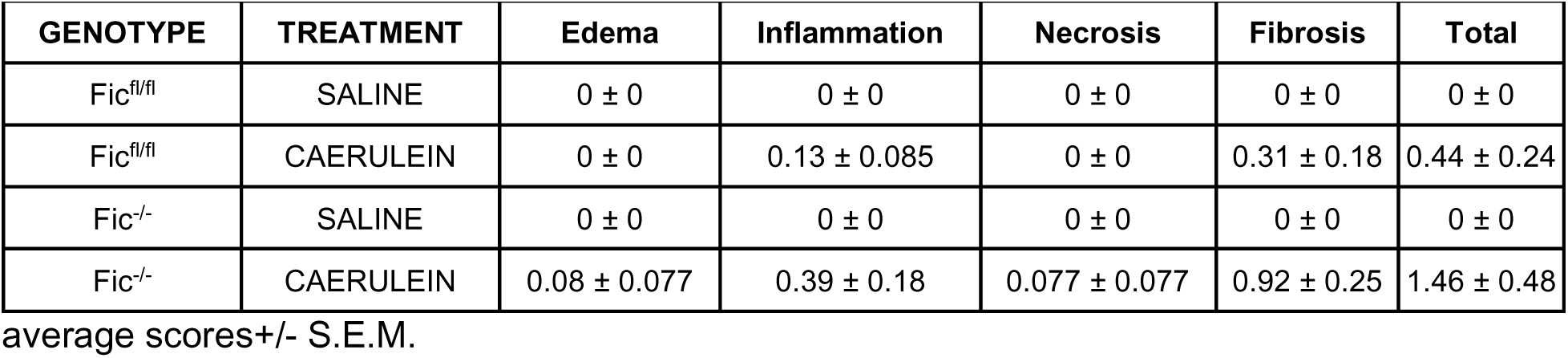
histopathological scoring of caerulein pancreatitis after 1 week recovery

## Discussion

The integrated stress response (ISR) plays a critical role in maintaining protein homeostasis during physiological stress and is triggered once the level of total intrinsic cellular stress reaches a level that may compromise the health of a cell. ISR responds to at least four cellular queues including ER-associated unfolded protein response (UPR), hypoxia, viral infection and nutrient deprivation. These pathways induce ISR by the inhibitory phosphorylation of the elongation translation initiation factor 2A (eIF2a), with levels of phosphor-eIF2a dictating the level of translation. When cellular stress levels reach maladaptive levels, recovery from stress becomes challenging due to saturating levels of phospho-eIF2a, stalled protein synthesis, and activation of apoptotic machinery. As UPR is one of the four pathways that trigger ISR, understanding regulatory mechanisms, such as posttranslational modifications of UPR machinery, are important for maintaining a high-fidelity response to cell and tissue stress.

To understand the role of reversible AMPylation of BiP in protein homeostasis in a mammalian system, we generated a conditional knockout of Fic in mice. We speculated that tissues reliant on secretion might be most affected by the deletion of Fic and therefore focused our initial efforts on the pancreas. Using both a fast-feeding and a caerulein-induced pancreatic injury model, we observed changes to UPR signaling and physiology of the pancreas suggestive of exocrine pancreas disfunction in *Fic^-/-^* mice. Analysis of UPR markers for these experiments revealed changes in the timing and duration of the UPR transcriptional response.

Furthermore, analysis of tissue recovery after light-induced or caerulein-induced damage in *Drosophila* eyes and mouse pancreas, respectively, indicates that loss of Fic reduces recovery from ER stress-associated tissue damage in both animal models. Therefore, the loss of Fic- mediated AMPylated BiP leaves tissues vulnerable to irreversible damage with chronic and repeated stresses.

Our and other groups have proposed that inactivating PTMs on BiP could provide a mechanism by which rapidly changing physiological fluctuations of the ER stress can be nimbly regulated (38). AMPylation of BiP by Fic allows for a pool of inactive chaperone to remain in the ER without deleterious consequences to protein folding, that might otherwise be hindered in the presence of excess chaperone (38, 39). Previous studies indicate that the pool of inactivated BiP is significant in various cell types, approximately 40% in fasted pancreas and over 50% in unstressed *Drosophila* S2 cells (17, 38). This inactive pool of BiP can then be readily activated to address increasing loads of unfolded proteins in cells with rapidly fluctuating demands on protein synthesis and secretion, such as the pancreas, while tempering the activation of the UPR.

We predict that Fic provides a necessary level of regulation of the UPR to properly adjust protein homeostasis in tissue with frequent physiological ER stress (**Figure 7**). By keeping a readily accessible pool of inactive BiP, cells can provide a nimble response to ER stress through a short burst of UPR activation. Rapid deAMPylation of BiP results in additional active chaperone much faster than what can be accomplished by new protein synthesis. This results in smaller, more moderate pulses of UPR signaling under repeated physiological stresses, keeping the cell in the beneficial *adaptive phase* of the UPR. As Fic is a UPR responsive gene, it’s possible as these stresses are repeated, a larger pool of inactive BiP may be generated over time, resulting in a more robust rapid response to unfolded proteins in the ER in these adapted tissues.

**Figure 7.**
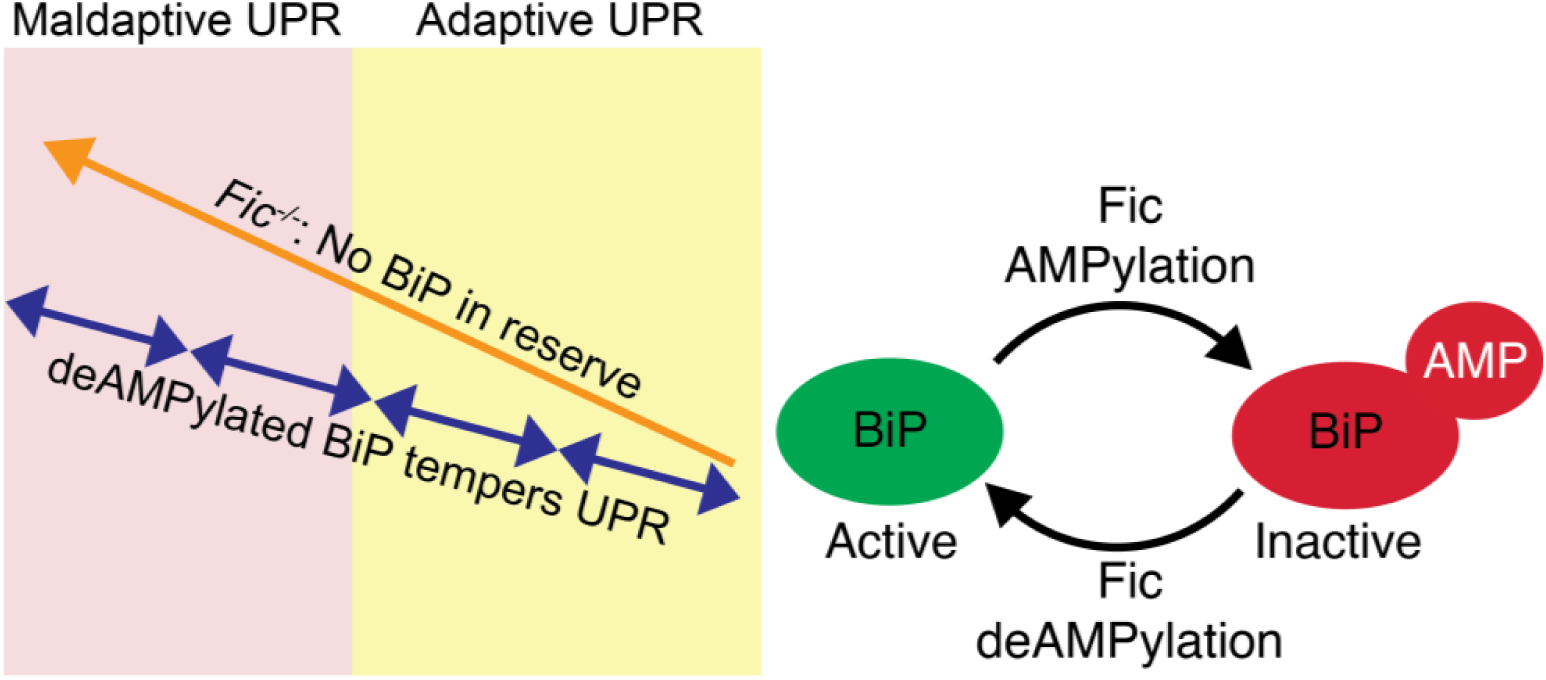
Model of Fic tempering of UPR.

Cells without Fic regulation of BiP lack this pool of chaperone on standby, resulting in prolonged and elevated UPR signaling as a delayed response requires more chaperone via transcription and translation to accommodate the increased physiological stress. Repetitive stress in *Fic^-/-^* tissues would result in amplified UPR, leading to progression into the *maladaptive phase* of the UPR and tissue damage over the lifetime of the tissue. Our data points to evidence of this in the exocrine pancreas, where elevated UPR signaling and serum Amylase indicate functional disruption. As elevated serum Amylase is one of the first key clinical indicators of pancreas dysfunction, we suspect additional and continued physiological stresses to *Fic^-/-^* tissue will lead to increased prevalence of disease.

The pancreas is primarily comprised of terminally differentiated professional secretory cells with limited regenerative capability. Therefore, the pancreas must employ mechanisms to ensure resilience to repetitive stress in order to last and properly function for the lifetime of the animal. Herein, we provide evidence that Fic provides one such mechanism through the moderation of the UPR during physiological stress. Similarly, a wild-type fly eye has the capacity to recover from the physiological stress of continuous light. In the absence of the Fic rheostat, the *fic^-/-^* eyes are challenged over time and lose the potential to regenerate rhabdomere integrity. Analogously, we observe more scarring in the injured *Fic^-/-^* pancreas.

Many studies to date have used tissue culture cell lines as a model to study the UPR in which a chemical stress is applied to cells resulting in a very strong, and frequently irreversible, induction of the UPR. Under these conditions, tissue culture cells respond in basically two ways: cell death or replication, allowing for new cells to overcome the stress. These options are far from optimal for differentiated cells within a tissue where cellular function needs to be maintained for survival of the organ and/or animal. We predict that many subtilties of UPR regulation will only be apparent under such physiological stresses in the context of specific tissues. Thus, it is not surprising that studies with tissue culture models have only exhibited very subtle differences in activation of UPR in the absence of Fic (20). Systems in an animal that use cells with high regenerative capacity and shorter lifespans may not require Fic mediation of the UPR as turnover and replenishment with new cells will bypass the need of rheostat. This is consistent with observations by other groups with a Fic deletion model (40). In sum, we predict that terminally differentiated post-mitotic cells will be principally reliant upon the Fic-mediated rheostat to maintain a healthy response to continuing physiologic stress over an animals lifetime.

### Ideas and Speculation

Whereas this study focuses on this one tissue only, we speculate that other tissues with professional secretory, terminally differentiated cells that must adapt to fluctuating stress will be similarly affected in the *Fic^-/-^* mouse. We propose the presence of Fic rheostat allows for tempering of the UPR response by maintaining a window for reversible UPR response that is critical for maintainance of protein homeostasis. The importance of this window has been highlighted with the treatment of UPR stress with the pharmacological agent ISRIB where it is only observed to be efficatious during the *adaptive phase* of UPR (9). Future studies with ISRIB and *Fic^-/-^* mice will be useful for understanding the importance of the Fic-mediated rheostat and treatment of disease.

For the health of an animal, it is critical to maintain resilence in terminally differentiated cells during repeated physiological stress to prevent disease. We predict that Fic regulation of the UPR will play a role in mitigating the deleterious effects of UPR activation in a variety of tissues with UPR-associated diseases, including retinal degeneration, athlerosclerosis, metabolic syndrome, and various neurodegenerative disorders. Our future studies will focus on the identification of tissues in which Fic plays a role in the regulation of the UPR and the physiological consequences of the absence of Fic-mediated regulation of the UPR.

## Materials and Methods

### Mice

The Institutional Animal Care and Use Committee of the University of Texas Southwestern Medical Center approved all experiments. Mice were housed in a specific pathogen-free facility and fed a standard chow diet (#2016 Harlan Teklad). C57BL/6J mice were purchased through the University of Texas Southwestern Mouse Breeding Core. For fasting studies, mice were separated into three groups fasted (for 16 hours), fasted-fed (fasted 14-hours, fed 2 hours), and fasted-fed-recovery (fasted 12 hours, fed 2 hours, fasted 2 hours). All mice were singly housed and fasted overnight (∼12-14 hours) with unrestricted access to water. After fasting, the fasted-fed group were provided unrestricted access to food for a 2 hour window and immediately sacrificed. The fasted-fed-recovery group provided unrestricted access to food for a 2 hour window and then food was removed for an additional 2 hours before sacrifice. Immediately upon sacrifice, blood was collected from each mouse and the pancreas was collected for analysis. Acute pancreatitis was induced as previously described (33) by administration of seven hourly intraperitoneal injections of caerulein (50 μg/kg) (Tocris Bioscience). Mice in the control group were injected with saline. For serum analysis, blood was collected in Microvette serum collection tubes (Sarstedt) and centrifuged at 10,000xg at 4°C for 15 minutes. Serum analysis was performed by *UT* Southwestern Metabolic Phenotyping Core using VITROS MicroSlide™ technology

### Generation of conditional Fic knockout

*Fic^fl/fl^* mice (C57Bl/6N background) harboring the conditional floxed Fic alleles were generated using CRISPR/Cas9 reagents at the Transgenic Technology Center of UT Southwestern Medical Center. The guide RNAs and donor ssODNs were designed to insert one *loxP* site upstream and another *loxP* site downstream of exon 3. The sgRNA sequences are 5’ ggggacctcccaatgtagag 3’ (upstream) and 5’ gctggcggttagggcctcac 3’ (downstream). Guides were selected using the CRISPR Design Tool (http://tools.genome-engineering.org). The crRNA and tracRNA were annealed and mixed with Cas9 protein to form a ribonucleotide protein complex. The donor ssODNs (IDT, Inc.) was added to the mixture and the cocktail was microinjected into the cytoplasm of fertilized, pronuclear staged eggs isolated from superovulated females. The eggs were incubated in media containing cytochalasin-B immediately before and during microinjection to improve egg survival. Alternatively, CRISPR reagents were delivered to the cytoplasm via electroporation using either a Nepa21 Super Electroporator (NEPAGENE, Ichikawa, Japan) or Gene Pulser (BioRad, Hercules, CA, USA). The surviving eggs were transferred into the oviducts of day 0.5 pseudopregnant recipient ICR females (Envigo, Inc.) to produce putative founder mice. Founder mice were identified via PCR using the primer set 5’ cagtgcagccatactgtagg 3’ and 5’ ccatgtggcttctgtgactc 3’ for the upstream *loxP* and primer set 5’ gccctggcccattacaaac 3’ and 5’ gctatcccctgccactcag 3’ for the downstream *loxP* site and the amplicons were submitted for Sanger sequencing. F0 mice were bred with C57Bl/6N or J mice to obtain F1 mice heterozygous for the floxed Fic allele (*Fic^+/f^*^l^), which were then intercrossed to produce *Fic^fl/fl^* mice. For deletion of *Fic*, *Fic^fl/fl^* mice were bred with C57Bl/6N mice carrying a *CAG-Cre* transgene(24). To confirm that *Fic* was deleted, 20 ng genomic DNA extracted from tail snips was used for genotyping by PCR with the following primers; upstream (5’ gggggtggttcaaggaag 3’ and downstream 5’ ctgcaacctcctactggc 3’ followed by restriction digestion with EcoR1 and BamH1).

### Histology

Mouse pancreas and liver were harvested and fixed in 10% neutral buffered formalin overnight at 4°C. Paraffin sections were embedded by the UT Southwestern (UTSW) Molecular Pathology Core. Samples were sectioned, and hematoxylin and eosin (H&E)–stained. H&E-stained pancreas was graded as previously described on an equal weight score (from 0 to 3) for edema (0 = absent; 1 = focally increased between lobules; 2 = diffusely increased; 3 = acini disrupted and separated), inflammatory infiltration (0 = absent; 1 = in ducts, around ductal margins; 2 = in the parenchyma, <50% of the lobules; 3 = in the parenchyma, >50% of the lobules), necrosis (0 = absent; 1 = periductal necrosis, <5% of cells; 2 = focal necrosis, 5 to 20% of cells; 3 = diffuse parenchymal necrosis, 20 to 50% of cells), and total severity score (sum of edema, inflammatory infiltrate, and necrosis scores). In addition H&E-stained pancreas from mice 1 week after caerulein treatment were graded for fibrosis (0 = absent; 1 = mild fibrosis between acini within at least one lobule, 2 = moderate fibrosis between acini with acinar drop-out in at least one lobule, 3 = severe fibrosis between acini with acinar drop-out in at least one lobule)

### Quantitative real-time PCR

RNA from pancreas and liver was extracted using RNA Stat-60 (Iso-Tex Diagnostics). Complementary DNA (cDNA) was generated from RNA (2 μg) using the High-Capacity cDNA Reverse Transcription Kit (Life Technologies). qPCR was performed by the SYBR Green method (41). Primer sequences for the genes analyzed can be found in **Table 3**. *U36B4* (<u>NM_007475</u>) was used as the reference mRNA. Experiments were performed on a BioRad CFX Touch and analyzed with CFX Maestro Software.

**Table 3.**
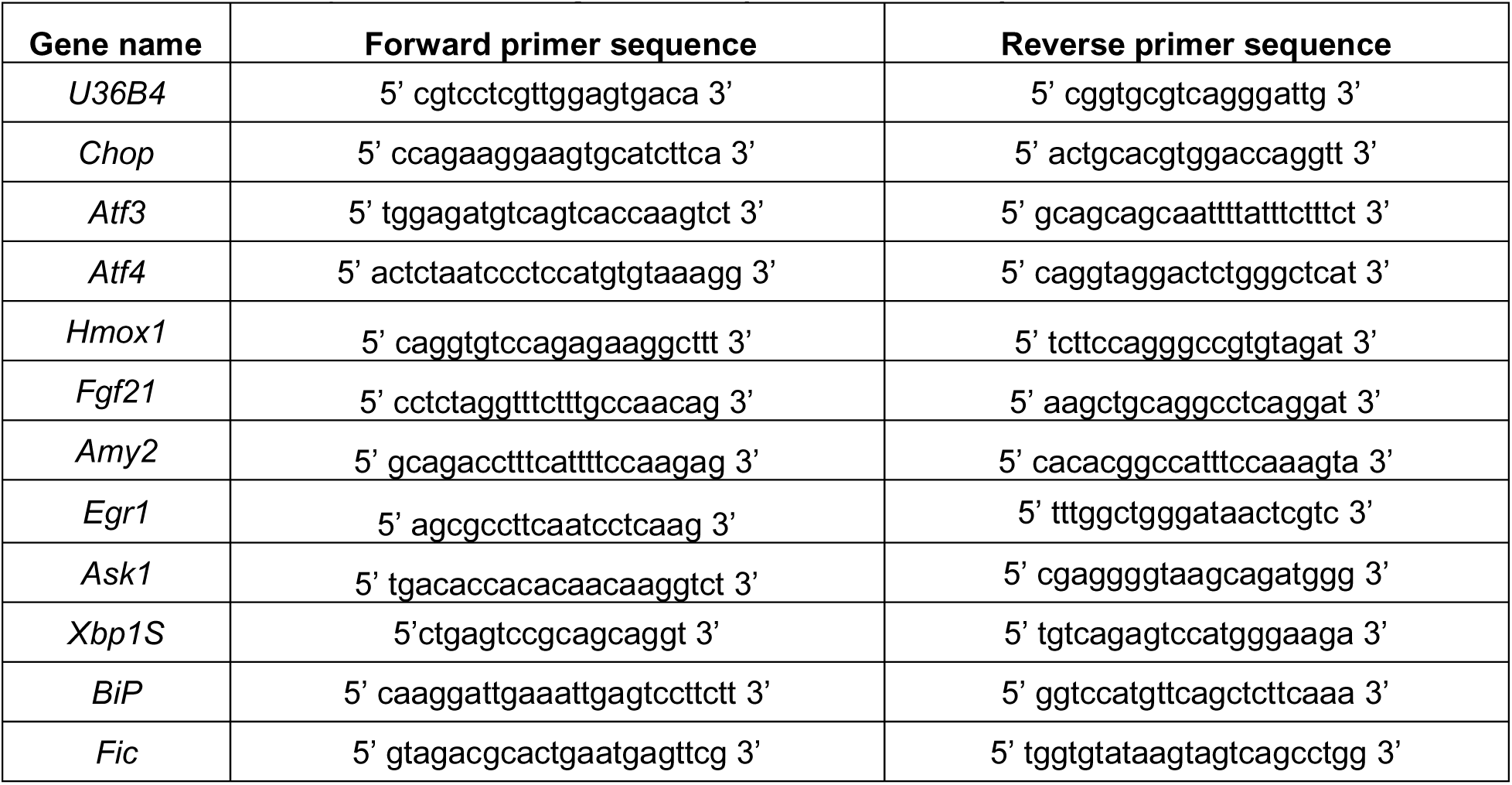
Primer sequences for the genes analyzed in this study

### Immunofluorescence of Brain Sections

Mouse brains were harvested and fixed in 10% neutral buffered formalin overnight at 4°C. Brains were equilibrated in 30% sucrose solution in 1x PBS and *embedded* with optimal cutting temperature (OCT) compound and frozen on dry ice. Samples were sectioned via cryostat at a thickness of 50μM. Sections were incubated for 1 hour in blocking buffer (5% normal serum, 1% BSA, 0.3% Triton X-100 in PBS), followed by overnight incubation at 4°C with primary antibodies anti-vimentin (Abcam) and anti-HYPE (custom antibody from Thermo/Fischer against GDVRPFIRFIAKCTET peptide) in blocking buffer. Sections were washed in PBS three times, followed by incubation for 1 hour with Alexa Fluor–conjugated secondary antibodies against the primary antibody’s host species in blocking buffer. Sections were washed in PBS three times, stained with 4′,6-diamidino-2-phenylindole (DAPI) (Life Technologies), and mounted with Aqua-Poly/Mount (Polysciences). Images were acquired with a Zeiss Axioscan.Z1 slide scanner provided by the Whole Brain Microscopy Facility at UTSW.

### Mass Spectrometry Analysis

Endogenous BiP samples from Fic^-/-^ and Fic^fl/fl^ mouse livers were run on SDS-PAGE gel and stained with Coomassie blue dye prior to analysis by mass spectrometry. Gel bands containing proteins were reduced with DTT for 1hr at 56°C and alkylated with iodoacetamide for 45min at room temperature in the dark. Samples were then digested with trypsin (MS grade) overnight at 37°C. Tryptic peptides were de-salted via solid phase extraction (SPE) prior to LC-MS/MS analysis. Experiments were performed on a Thermo Scientific EASY-nLC liquid chromatography system coupled to a Thermo Scientific Orbitrap Fusion Lumos mass spectrometer. To generate MS/MS spectra, MS1 spectra were first acquired in the Orbitrap mass analyzer (resolution 120,000). Peptide precursor ions were isolated and fragmented using high-energy collision-induced dissociation (HCD). The resulting MS/MS fragmentation spectra were acquired in the ion trap. MS/MS spectral data was searched using the search engine Mascot (Matrix Science). Precursor and product ion mass tolerances were set to 15 ppm and 0.6 Da, respectively. Three missed cleavages were allowed. Modifications included carbamidomethylation of cysteine (+57.021 Da), oxidation of methionine (+15.995 Da), and AMPylation of serine/threonine/tyrosine (+329.053 Da). MS/MS spectra of AMPylated peptides were manually searched and verified. Biological replicates of each sample were analyzed.

### Western Blot Analysis

Pancreas was washed in PBS and homogenized in isolation buffer (20mM Tris pH 7.4, 5mM NaF, 1mM EDTA, 1mM EGTA, 250mM Sucrose, PMSF, PhosSTOP (Roche), Protease Inhibitor Cocktail (Roche)) with a Teflon-coated dounce homogenizer. Lysates were centrifuged at 600xg for 10 mintues three times at 4°C to remove nuclei and cellular debris (P1). The supernatant (S1) was collected and centrifuged at 13,000xg for 15 minutes to pellet ER and mitochondrial membranes (P2) for western blot analysis. Supernatant (S2) was reserved for western blot analysis. Lysates were separated by SDS-PAGE and transferred to PVDF membranes. Phos tag analysis was performed as previously described (29).Blots were probed with anti-AMP 17g6 (gift from Aymelt Itzen), anti-GRP78 (Abcam), anti-tubulin (Abcam), anti-eif2A (Cell Signaling), anti- EIF2S1-phospho (Abcam), anti-IRE1 (Cell Signaling), and anti-IRE1-phospho (Novis). Membranes were then incubated with horseradish peroxidase–conjugated secondary antibodies (Cell Signaling Technology) against the primary antibody’s host species for 1 hour. Membranes were developed using the ECL substrate solution (Bio-Rad). Quantification of western blots was performed using NIH ImageJ software. Band densities were measured and subtracted from background.

### Fly stocks and rearing conditions

Bloomington Stock Center provided *w*^1118^ (BS# 3605). The *fic*^30C^ flies was previously described (21). All flies were reared on standard molasses fly food, under room temperature conditions. For light treatments, flies were collected within one to two days of enclosing, and placed in 5cm diameter vials containing normal food, with no more than 25 flies, and placed at either LD (lights ON 8am/lights OFF 8pm) or LL. The same intensity white LED light source was used for both conditions and flies were kept the same distance away from the light source, which amounted to approximately 500 lux. LD and LL treatments were done at 25°C.

### Deep pseudopupil analysis

Flies were anesthetized on CO_2_ and aligned with one eye facing up. Using a stereoscopic dissection microscope, each fly was scored for presence or loss of the deep pseudopupil (42), and the percentage of flies with intact pseudopupils was calculated. For each genotype/treatment, over 50 flies were scored per replica and three biological replicas were performed (n=3).

## Author contributions

A.K.C., G.H. and K.O. designed the experiments involving mice. A.M. and H.K. designed and conducted the experiments involving flies. A.K.C., H.F.G, G.H., N.S, B.G., K.S. and A.B., conducted experiments involving mice. K.A.S. performed mass spectrometry analysis on prepared samples. H.F.G. and S.C. designed conditional null allele of Fic. H.A.F. performed histological sectioning and staining of tissue. B.E. and E.D performed histopathological analysis of stained tissue. A.K.C. and K.O. wrote the manuscript with input from all authors.

## Acknowledgments

We thank Robert Hammer and the Transgenic Technology Center of UT Southwestern Medical Center for their assistance in generation of the *Fic^fl^* allele in the C57Bl/6N mouse. Thank you to Eric Olson and Rhonda Bassel-Duby for their gift of CAG-Cre transgenic mice. We want to thank UT Southwestern Metabolic Phenotyping Core for the analysis of serum metabolites (Amylase, Lipase, and AST) and expertise and UT Southwestern Whole Brain Microscopy Facility at UTSW for use of their instruments and guidance. We thank Aymelt Itzen for his generous gift of monoclonal anti-AMP antibodies. We thank the Orth lab members for discussions and editing and Ricardo Rayas, Alexander LaFrance, Nalleli Rodrigez for assistance with handling of mice.

