## Supplementary material for "Fic-mediated AMPylation tempers the Unfolded Protein Response during physiological stress": Supp File

```
FicD_Flag     MILMPMASVVAVAEPKWVSVWGRFLWMALLSMALGSLLALLLPLGVVEEHCLAVLRGFHL     60
FicD_WT       MILMPMASVVAVAEPKWVSVWGRFLWMALLSMALGSLLALLLPLGVVEEHCLAVLRGFHL     60
FicD_Null     MILMPMASVVAVAEPKWVSVWGRFLWMALLSMALGSLLALLLPLGVVEEHCLAVLRGFHL     60
              ************************************************************

FicD_Flag     LRSKLDRAQPVVPKCTSLCTELSVSSRDAGLLTVKTTASPAGKLEAKAALNQALEMKRQG     120
FicD_WT       LRSKLDRAQPVVPKCTSLCTELSVSSRDAGLLTVKTTASPAGKLEAKAALNQALEMKRQG     120
FicD_Null     LRSKLDRAQPVVPKCTSLCTELSVSSRDAGLLTVKTTASP--------------------     100
              ****************************************

FicD_Flag     KRGKAHKLFLHALKMDPGFVDALNEFGIFSEEDKDIIQADYLYTRALTISPFHEKALVNR     180
FicD_WT       KRGKAHKLFLHALKMDPGFVDALNEFGIFSEEDKDIIQADYLYTRALTISPFHEKALVNR     180
FicD_Null     ------------------------------------------------------------     100

FicD_Flag     DRTLPLVEEIDQRYFSVIDSKVKKVMSIPKGSSALRRVMEETYYHHIYHTVAIEGNTLTL     240
FicD_WT       DRTLPLVEEIDQRYFSVIDSKVKKVMSIPKGSSALRRVMEETYYHHIYHTVAIEGNTLTL     240
FicD_Null     ------------------------------------------------------------     100

FicD_Flag     SEIRHILETRYAVPGKSLEEQNEVIGMHAAMKYINTTLVSRIGSVTMDDMLEIHRRVLGY     300
FicD_WT       SEIRHILETRYAVPGKSLEEQNEVIGMHAAMKYINTTLVSRIGSVTMDDMLEIHRRVLGY     300
FicD_Null     ------------------------------------------------------------     100

FicD_Flag     VDPVEAGRFRRTQVLVGHHIPPHPRDVEKQMQEFTQWLNSEDAMNLHPVEFAALAHYKLV     360
FicD_WT       VDPVEAGRFRRTQVLVGHHIPPHPRDVEKQMQEFTQWLNSEDAMNLHPVEFAALAHYKLV     360
FicD_Null     ------------------------------------------------------------     100

FicD_Flag     YIHPFIDGNGRTSRLLMNLILMQAGYPPITIRKEQRSEYYHVLEVANEGDVRPFIRFIAK     420
FicD_WT       YIHPFIDGNGRTSRLLMNLILMQAGYPPITIRKEQRSEYYHVLEVANEGDVRPFIRFIAK     420
FicD_Null     ------------------------------------------------------------     100

FicD_Flag     CTEVTLDTLLLATTEYSVALPEAQPNHSGFKETLPVDYKDHDGF*    464
FicD_WT       CTEVTLDTLLLATTEYSVALPEAQPNHSGFKETLPVRP*------    458
FicD_Null     ---------------------------------------------    100
```

**Figure 1 – Supplement 1** Multiple sequence alignment of Fic. The predicted protein sequences of *Fic*$^{fl}$ (FicD_FLAG), *Fic* (FicD_WT), and *Fic*$^{-}$ (FicD_Null) were aligned. Fic$^{fl}$ sequence is identical to Fic with the addition of a 6 amino acid FLAG sequence on the C-terminus of the protein. Fic$^{-}$ sequence results in a truncated 100 amino acid protein.

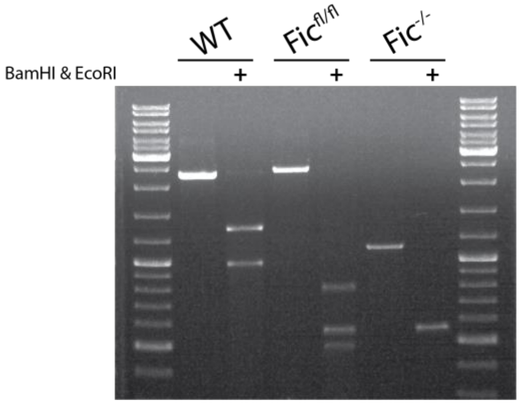


**Figure 1 – Supplement 2** PCR genotyping of Fic alleles. Representative agarose gel of PCR amplicons and digests. Primers were used that amply across the modified region of Fic, producing a 2306bp $Fic^{+}$ amplicon, 2407bp $Fic^{fl}$ amplicon, and 1118bp $Fic^{-}$ amplicon. Digestion of Fic amplicons with BamHI and EcoRI result in DNA fragments of 1318 and 988 bp ($Fic^{+}$), 802, 565, 556, and 484 ($Fic^{fl}$), and 556 and 562 ($Fic^{-}$).

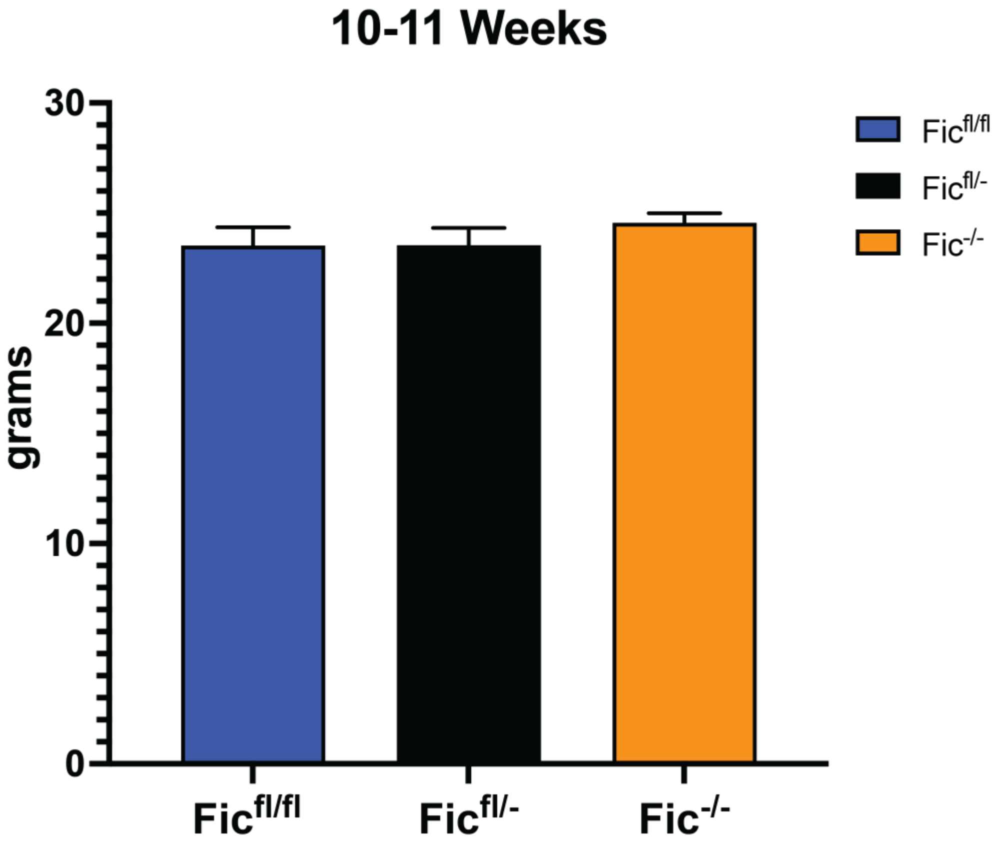


**Figure 1 – Supplement 3.** 10-11 week old male *Fic*$^{fl/fl}$ (N= 8), *Fic*$^{fl/-}$ (N=9), and *Fic*$^{-/-}$ (N=9) littermates were weighed before fasting. Graph represent weight of mice. Bars indicate mean weight and error bars represent standard error.

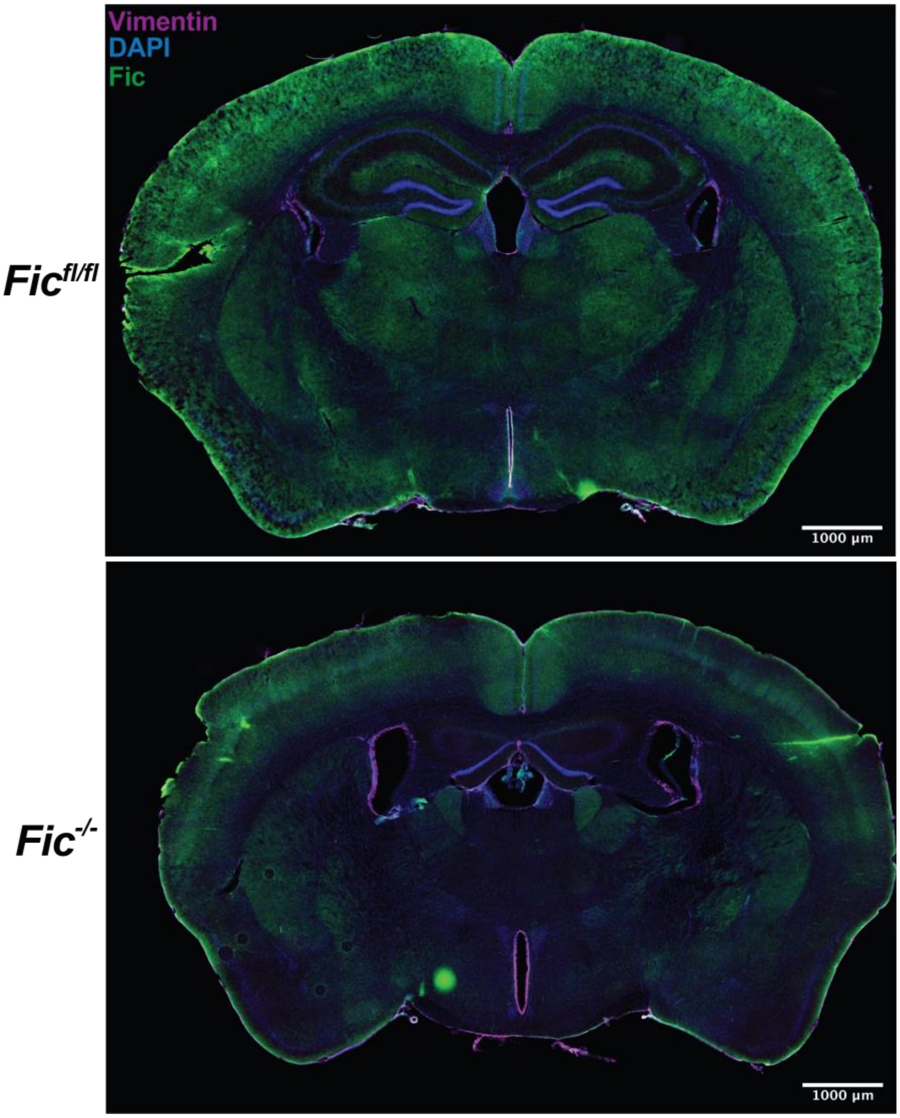


**Figure 1 – Supplement 4.** Representative image of anti-Fic and anti-Vimentin immunohistochemistry in coronal section of murine brain. Scale bar, 1000μM.

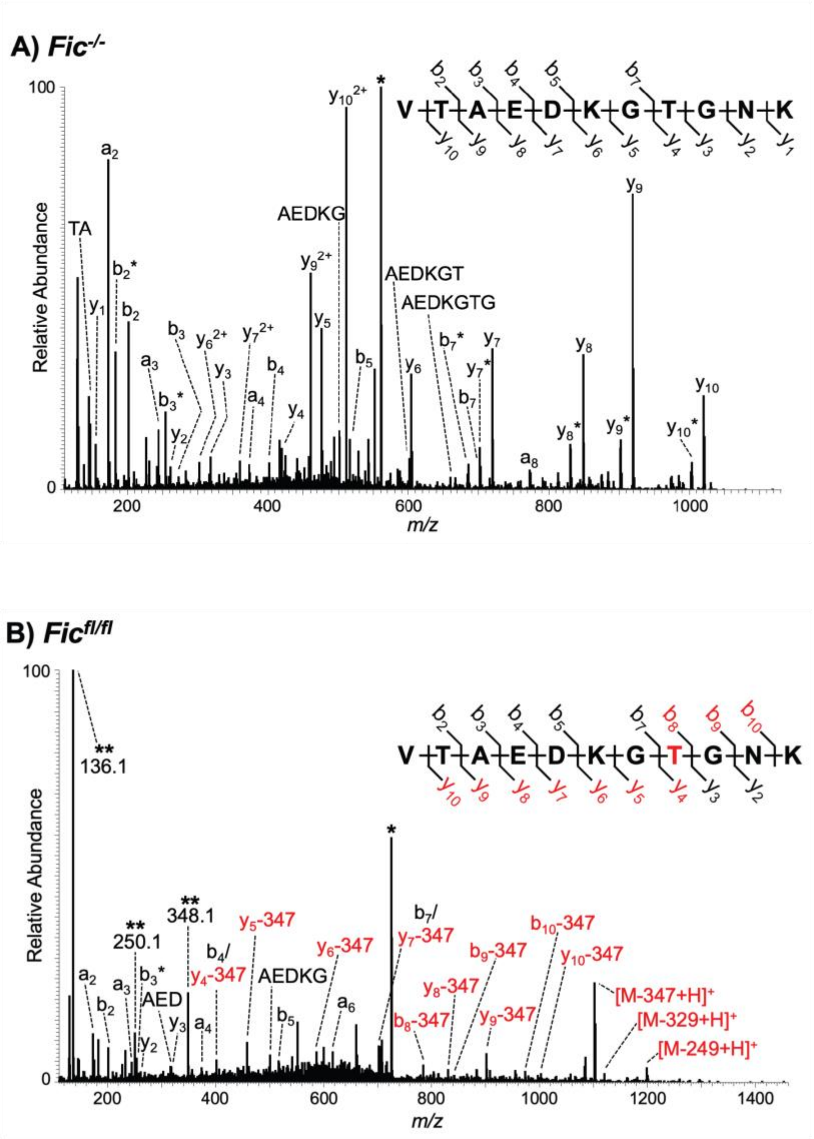


**Figure 1 – Supplement 5.** MS/MS spectra of **A)** unmodified peptide ion VTAEDKGTGNK from Fic$^{-/-}$ mouse and **B)** AMPylated peptide ion VTAEDKGTGNK from Fic$^{fl/fl}$ mouse. The AMPylated threonine residue is highlighted in red while AMPylation was not observed in Fic$^{-/-}$. The precursor $[M+2H]^{2+}$ ion is labeled with a single asterisk (*) and spectra were generated via HCD fragmentation. Fragment ions containing the AMPylated threonine residue (red) show characteristic mass shifts corresponding to loss of the AMP group (-347 Da). Unique ions corresponding to neutral loss of the AMP group (labeled with **) are also present at 136.1, 250.1, and 348.1 Da. B- and y- fragment ions labeled with a single asterisk (*) correspond to neutral loss of $H_2O$ (-18 Da).

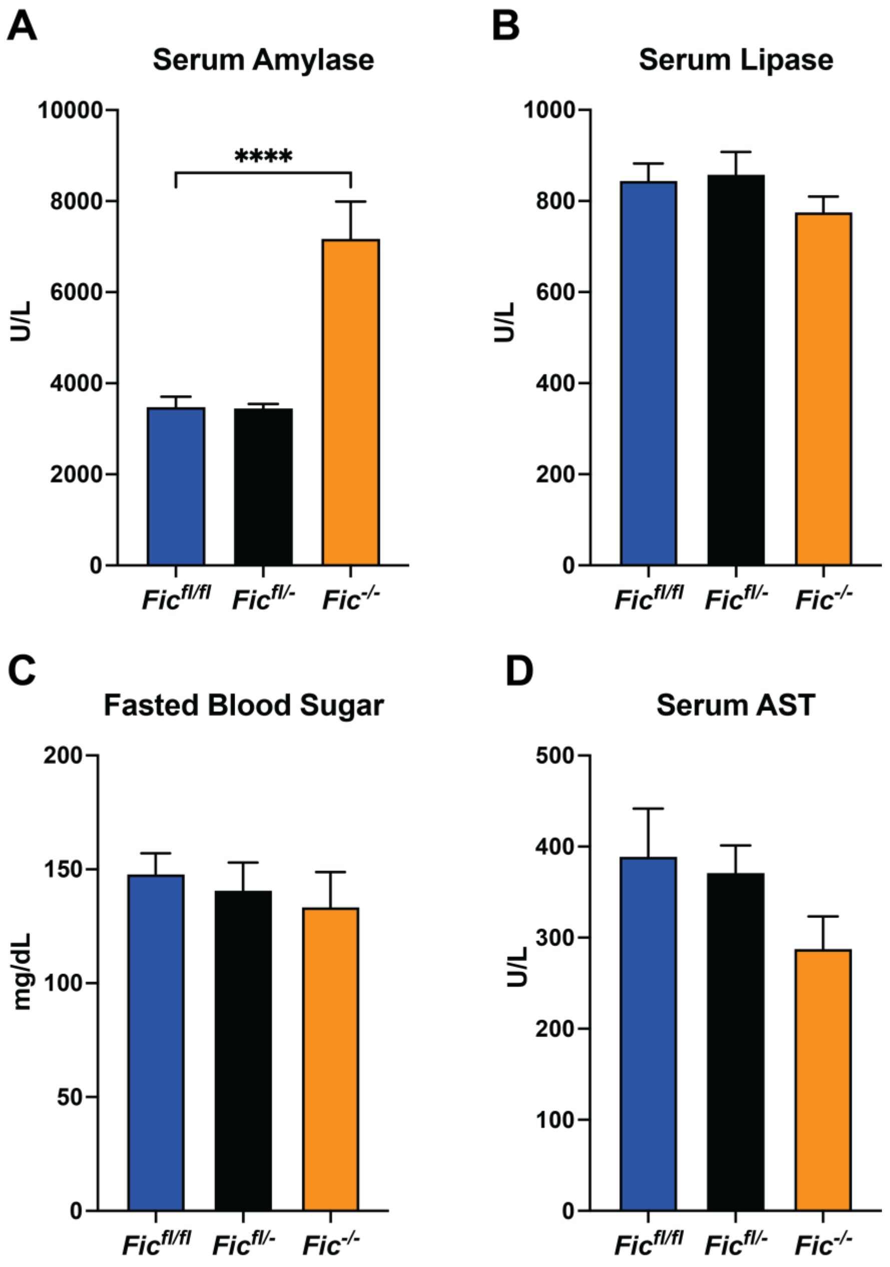


**Figure 2 – Supplement 1.** A) Quantification of serum Amylase in *Fic*$^{fl/fl}$*, Fic*$^{fl/-}$*, and Fic*$^{-/-}$ mice under fasted condions. B) Quantification of serum Amylase in *Fic*$^{fl/fl}$ *and Fic*$^{-/-}$ mice under fasted condions. C) Quantification of blood glucose in *Fic*$^{fl/fl}$ *and Fic*$^{-/-}$ mice under fasted condions. D) Quantification of serum aspartate aminotransferase (AST) in *Fic*$^{fl/fl}$ *and Fic*$^{-/-}$ mice under fasted condions. Bars indicate mean and error bars represent standard error. Statistics were performed using GraphPad Prism 9 using an 1-way ANOVA. N=8-9. ****, $p < 0.0001$.

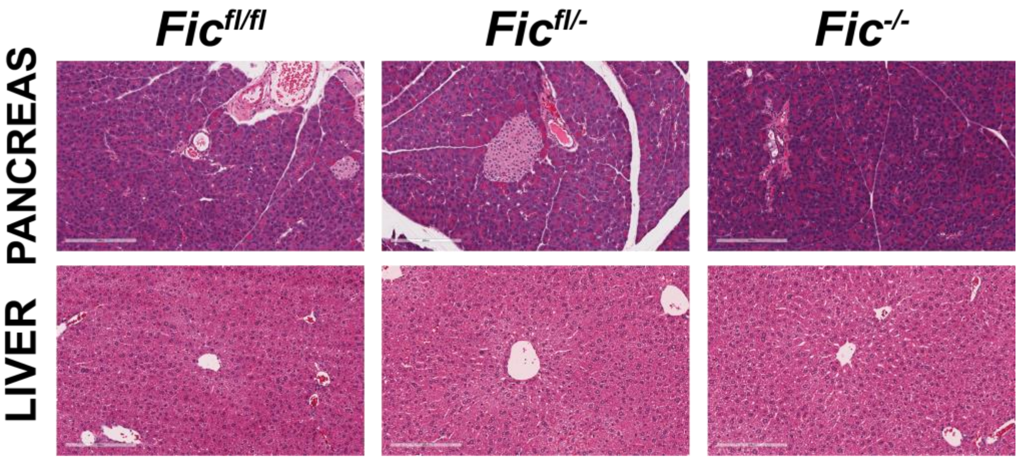


**Figure 2 – Supplement 2.** Representative hematoxylin and eosin stained images of pancreas and liver from *Fic$^{fl/fl}$, Fic$^{fl/-}$, and Fic$^{-/-}$* mice. Scale bar, 200μM

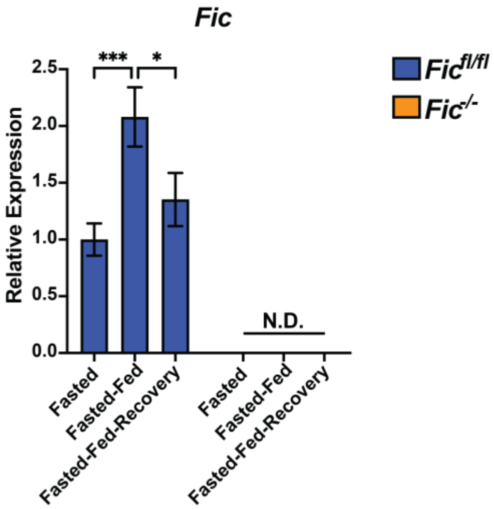


**Figure 3 – Supplement 1.** Quantification of *Fic* mRNA analyzed by qPCR from *Fic*$^{fl/fl}$ (blue bar) and *Fic*$^{-/-}$ (orange bar) mouse pancreas after fasting, fast-feeding, and fast-feed-recovery. Expression values were normalized to that of the housekeeping gene *U36B4.* Bars indicate mean relative expression compared to fasted controls, and error bars represent standard error. Fic mRNA was below detection cutoff in *Fic*$^{-/-}$ samples. Statistics were performed using GraphPad Prism 9 using an 2-way ANOVA. N=8. N.D., not detected; *, $p < 0.05$; ***, $p < 0.001$.

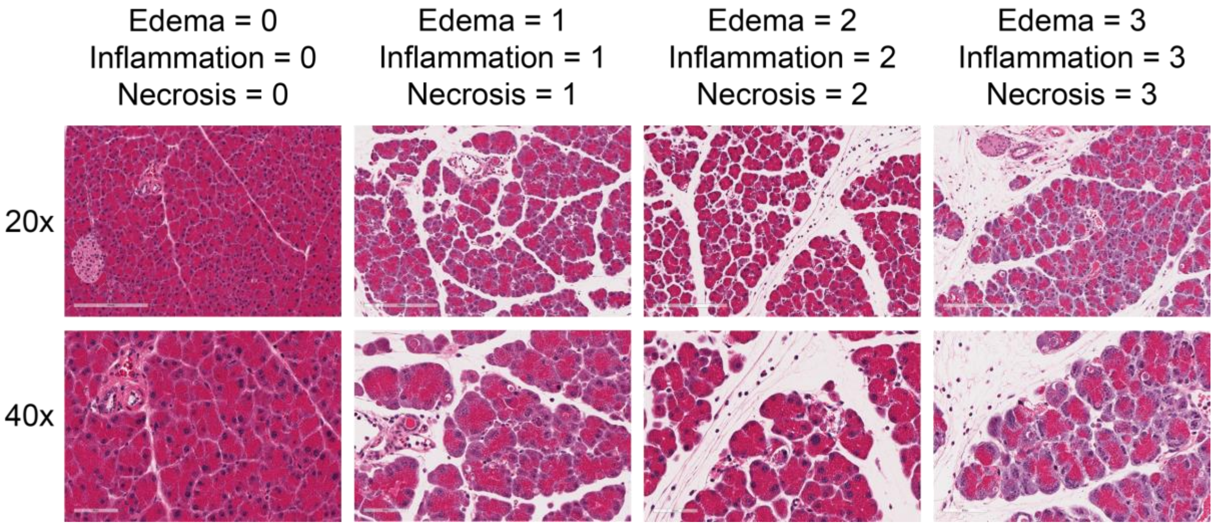


**Figure 4 – Supplement 1.** Representative hematoxylin and eosin stained images of pancreas with severity of pancreatic edema, inflammatory infiltrate, necrosis of 0, 1, 2 and 3. Images at 20x (scale bar, 200μM) and 40x (scale bar, 60μM) shown.

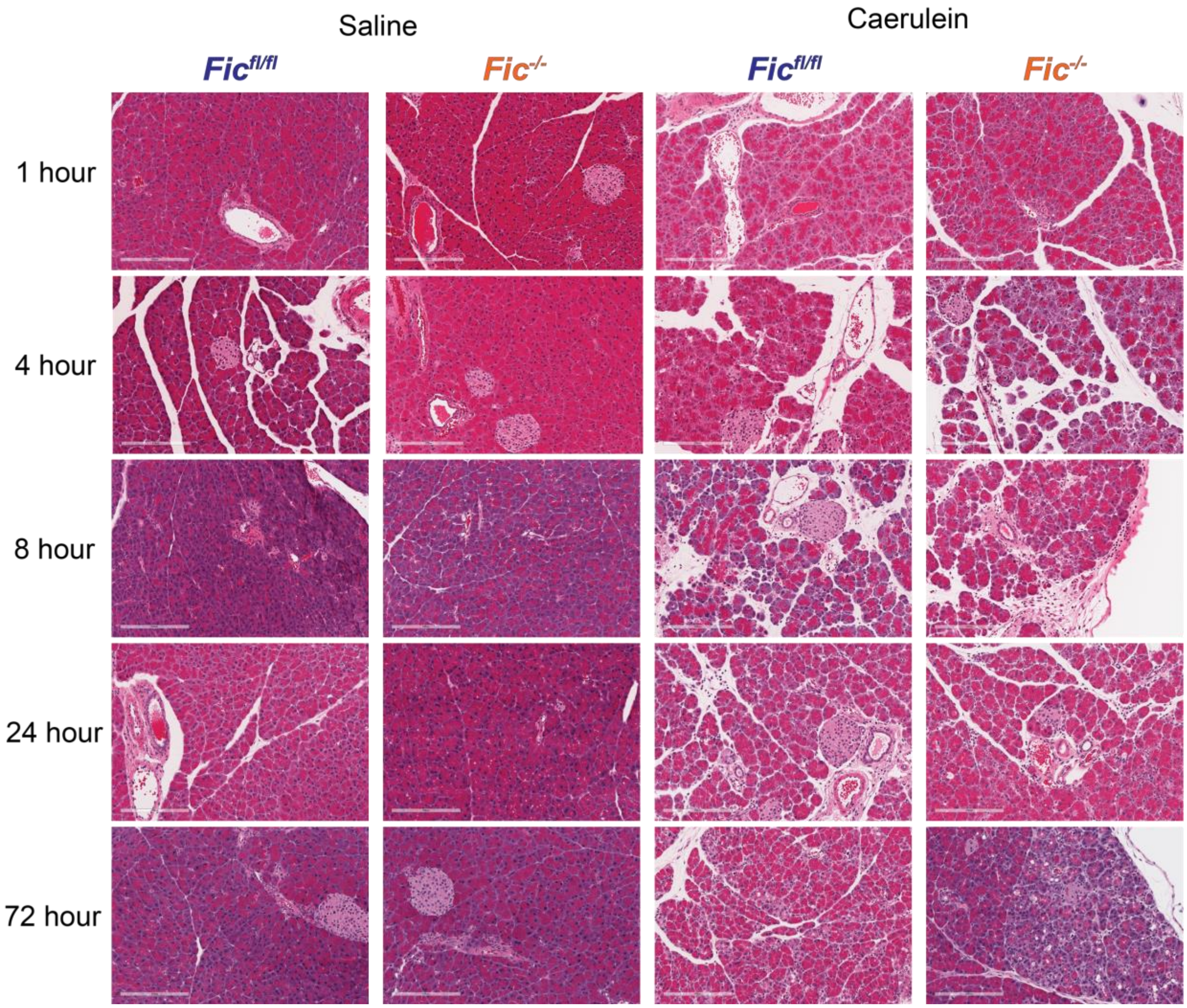


**Figure 4 – Supplement 2.** Representative hematoxylin and eosin stained images of pancreas of *Fic*$^{fl/fl}$ and *Fic*$^{-/-}$ mice treated with saline or caerulein at 1, 4, 8, 24, and 72 hours after first injection. Scale bar, 200μM.

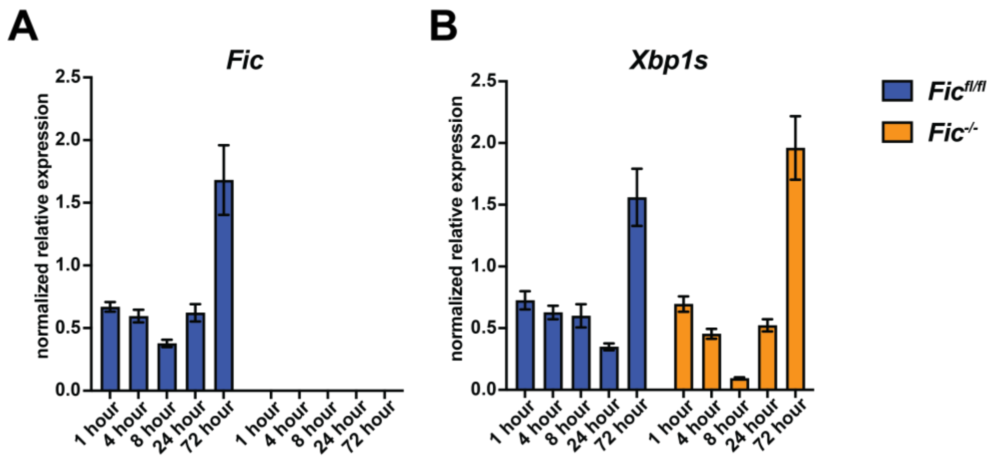


**Figure 5 – Supplement 1.** Quantification of A) *Fic* and B) *Xbp1* mRNA analyzed by qPCR from *Fic*$^{fl/fl}$ (blue bar) and *Fic*$^{-/-}$ (orange bar) mouse pancreas during 72 hours of caerulein-induced acute pancreatitis and recovery. Expression values were normalized to that of the housekeeping gene *U36B4.* Bars indicate mean relative expression compared to Saline-treated controls at each timepoint. N=5-7.
